# Sequestration-based Protein Neural Networks Tolerate the Effects of Shared Translational Resources

**DOI:** 10.64898/2025.12.01.691641

**Authors:** Enhao He, Frank Britto Bisso, Christian Cuba Samaniego

**Affiliations:** Ray and Stephanie Lane Computational Biology Department, Carnegie Mellon University, Pittsburgh, PA, USA; Joint CMU-PITT PhD Program in Computational Biology, Pittsburgh, PA, USA

## Abstract

Biochemical neural networks offer a promising framework for implementing advanced computation in living cells, yet how competition for shared cellular resources shapes their behavior remains poorly characterized. In this work, we develop a resource-aware mathematical framework to analyze how competition for ribosomes affects protein neural networks implemented via molecular sequestration in bacterial systems. Focusing on classification tasks, we show that when all mRNA species share identical ribosome-binding affinities, competition induces a uniform rescaling of the network’s effective weights, reducing the output magnitude while preserving the geometry of the decision boundary. When these affinities differ, competition introduces curvature in the decision boundary, transforming a linear classifier into a nonlinear one through resource coupling. This curvature is constrained to a well-defined region of the input space whose extent is determined by the degree of variability in the binding affinities, making the effect both predictable and, potentially, exploitable for design. Next, we numerically extend our analysis from a single perceptron to multi-layer architectures with two hidden-layer nodes and show that, for both closed and non-convex decision boundaries, resource competition preserves the boundary geometry in the homogeneous-affinity case, while in the heterogeneous case, curvature distortions from hidden-layer nodes propagate and compose at the output within a bounded uncertainty region. Finally, we incorporate growth-mediated feedback into the framework and show that the structural form of the activation function and decision boundary is preserved. Overall, our results provide a quantitative basis for understanding how translational resource competition shapes the behavior of protein-level molecular classifiers.

## I. INTRODUCTION

Cells have evolved to process a wide range of input combinations and reliably integrate them through signaling pathways to produce specific phenotypes, as exemplified by the immune system [1] and developmental programs [2, 3]. Replicating this level of information processing in synthetic circuits, however, remains a major challenge. One promising approach is to leverage the concept of neural networks, which provides a quantitative framework for designing circuits that integrate multiple inputs and produce discrete outputs. In classification tasks, these networks implement decision-making through a *decision boundary*, whereby combinations of input signals on one side of the boundary trigger one cellular response, while inputs on the other side trigger a different response [4]. Existing implementations of such networks have primarily been realized *in vitro* using DNA strand-displacement reactions [5–8]. More recently, a small number of studies have demonstrated related designs in living cells [9–11]. However, how competition for finite cellular resources affects classifier performance, and in particular the geometry and location of the decision boundary, has not been systematically analyzed.

Resource competition arises when synthetic and endogenous genetic circuits draw on the same molecular machinery, such as ribosomes, RNA polymerases, or other shared components. Under these conditions, it cannot be assumed that the behavior of a composed genetic circuit follows from the characterization of its individual parts, since shared resource usage introduces indirect coupling between otherwise independent modules. In general, there are two common situations that illustrate how this loss of composability emerges in practice. Changes in the cellular environment, such as shifts in pH, carbon sources, or temperature, trigger adaptive responses that reallocate cellular resources, reducing those available to the synthetic circuit and thereby affecting its function [12, 13]. Conversely, enforcing high levels of synthetic gene expression diverts resources away from essential cellular processes, often leading to reduced growth [14, 15]. This growth-mediated feedback in turn alters global transcriptional and translational capacities, perturbing circuit function. In this work, we focus on bacterial systems, where ribosomes constitute the primary limiting resource [16, 17] and where competition for translational capacity has been most extensively characterized experimentally [18, 19].

In this setting, designing reliable molecular classifiers requires explicitly accounting for how shared translational resources affect the decision boundary. Here, we develop a quantitative framework to analyze proteinbased neural networks operating under competition for ribosomes. Within this general setting, we focus on sequestration-based implementations in which an output protein reversibly binds to another protein to form an inactive complex, and the production of both proteins is subject to competition for the translational machinery. Using this framework, we quantify how the endogenous demand for ribosomes impacts the behavior of both linear and nonlinear classifiers, and we further extend the framework to incorporate growth-mediated feedback.

## II. RESULTS

Throughout the manuscript, we denote species with uppercase letters (e.g., *Y*) and their concentrations with the corresponding lowercase letters (e.g., *y*).

### A. Quantifying the effect of shared translational resources in protein production

To understand how protein production is affected by shared translational resources, we analyze the translation mechanism by modeling the competition of messenger RNA (mRNA) molecules for the available pool of ribosomes (*R*). We consider the *i*-th input species *X*_*i*_ producing mRNA at a rate constant *α*_*i*_. This mRNA (*M*_*i*_) can reversibly bind to a ribosome to form a complex (*R*_*i*_) at rate constant *a*_*i*_, which dissociates at rate *d*_*i*_. This complex enables translation into a protein (*Y*_*i*_) at rate *v*_*i*_. Finally, we consider that all species decay due to dilution at rate *δ*, and explicitly model the degradation of both free mRNA and ribosome-bound mRNA with the rate constant *ϕ*. We index the mRNA species as *i* = 1, … , *n*_2_, where *i* = 1, … , *n*_1_ corresponds to transcripts from the synthetic circuit and *i* = *n*_1_ + 1, … , *n*_2_ corresponds to endogenous transcripts. This indexing defines the following set of chemical reactions:

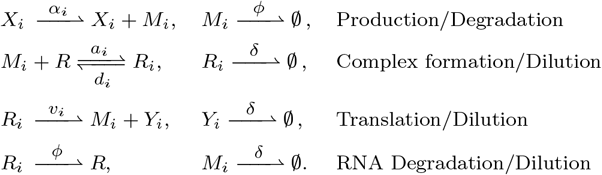

Using the law of mass action, we can model these chemical reactions with the following set of Ordinary Differential Equations (ODEs):

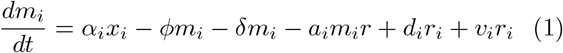

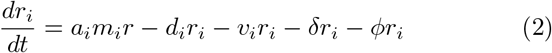

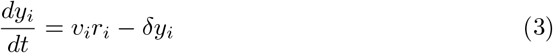

We can further quantify the effect of resource competition by defining a resource demand term for each mRNA that captures the fraction of the finite ribosome pool allocated to its translation, similar to resource demand coefficients used to model competition effects in genetic circuits [13, 20]. From the steady-state analysis detailed in

Appendix A, we can rewrite the equation of mass conservation for the ribosome pool as indicated by the following expression:

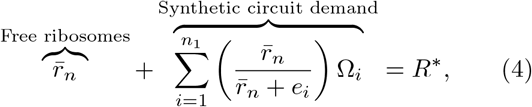

where 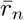 is the normalized steady-state level of free ribosomes, *e*_*i*_ is the normalized binding affinity between the *i*-th mRNA and a ribosome (or, equivalently, the dissociation constant of the *i*-th mRNA-ribosome complex), and Ω_*i*_ is the effective production rate of protein *Y*_*i*_, with all quantities normalized by the total ribosome pool (*r*^tot^). The variable *R*^*\**^ represents the effective ribosome availability after accounting for endogenous demand, and can be further defined as

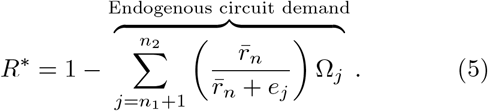

Both Eqs. (4) and (5) describe how the total ribosome pool is divided between free ribosomes and those engaged in translation at steady-state. Ribosomes bound during translation can be separated into those engaged by endogenous transcripts, corresponding to a fraction 1 − *R*^*\**^, and those available to the synthetic circuit, given by *R*^*\**^. Moreover, for the *i*-th synthetic transcript, we define the ribosome allocation factor (*S*_*i*_) as follows:

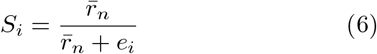

Then, the value *S*_*i*_Ω_*i*_ describes the effective protein *Y*_*i*_ production rate, showing how free ribosome availability and the mRNA–ribosome binding affinity (*e*_*i*_) constrain translation. In the following sections, we examine how variations in *R*^*\**^ and binding affinities influence overall synthetic circuit behavior.

### B. Translational resource competition scales the output but maintains the thresholding behavior in a protein-level neural network

To evaluate how shared translational resources influence the behavior of a molecular perceptron, we build on previous work [9, 21] and adopt a sequestration-based circuit as our synthetic system. The full chemical reaction network is shown in Fig. 1-A, while the abstraction using the conventional notation for a perceptron – the fundamental unit of a feedforward neural network – is shown in Fig. 1-B. Briefly, the circuit is composed of two input species, *X*_1_ and *X*_2_, that drive transcription of their corresponding mRNAs (*M*_1_ and *M*_2_), which compete for binding to the ribosome pool *R*. Each resulting complex produces the proteins *Y*_1_ and *Y*_2_, respectively. These proteins reversibly bind to one another at rate *b* to form a complex *C*, which dissociates at rate *u* and is diluted at rate *δ*. To model within the framework of Eqs. (1)-(3), we restrict the index to *i* = {1, 2, 3} . The species *i* = {1, 2} correspond to the molecular sequestration reaction, modeled as:

**FIG. 1.**
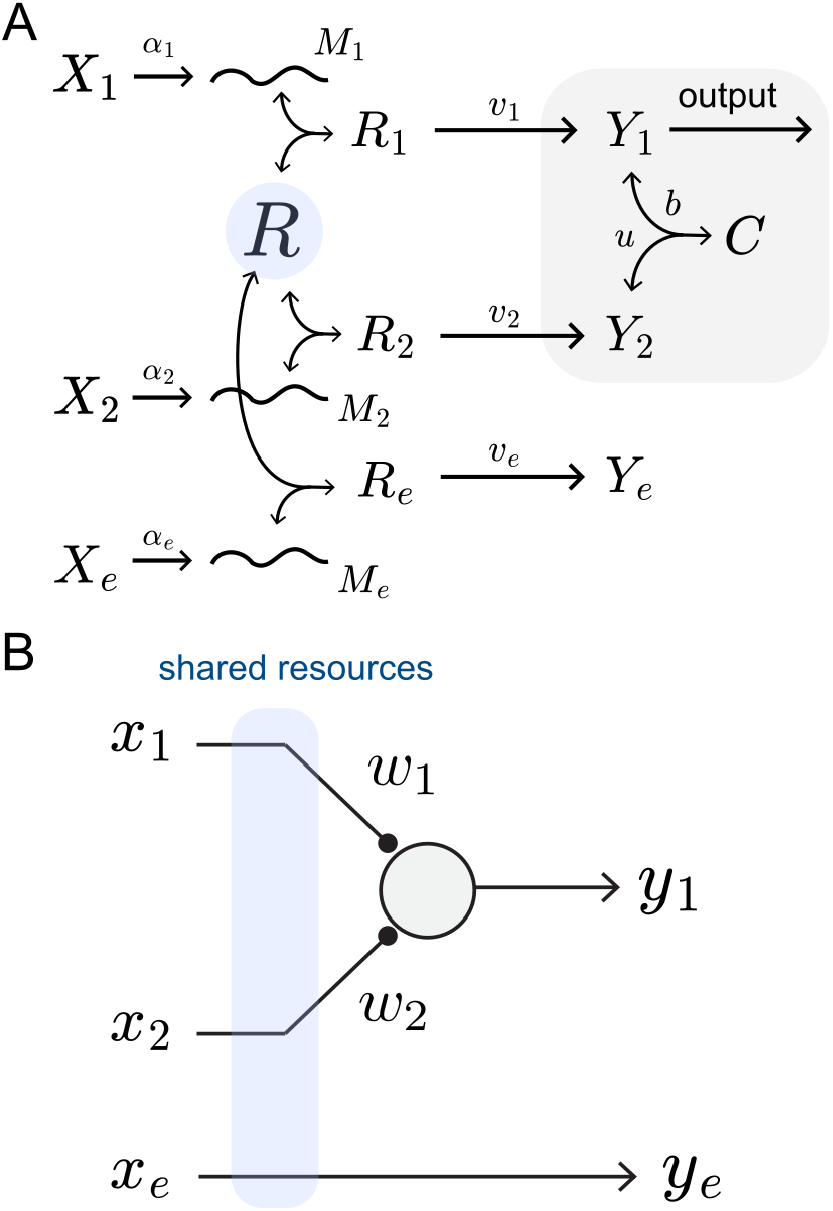
Sequestration-based perceptron under ribosomal competition. (A) Schematic of the chemical reaction network showing the sequestration mechanism (grey) and competition among mRNAs (*M*_1_ and *M*_2_) and endogenous transcripts (*M*_*e*_) for ribosomes (*R*). (B) Simplified representation highlighting competition for the ribosome pool, indicated in blue.xs

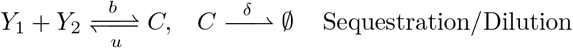

Using the law of mass action, we obtain the ODEs that model the sequestration-based perceptron:

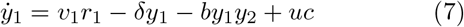

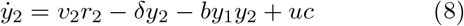

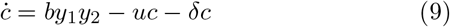

The remaining index, *i* = 3, corresponds to an additional input *X*_*e*_ that produces the transcript *M*_*e*_, which gets translated into the *Y*_*e*_ protein. The species *M*_*e*_ competes for the same ribosome pool and is used as a minimal model of the endogenous translational demand.

From the steady-state analysis detailed in Appendix B, the output of the sequestration-based perceptron can be expressed in the fast sequestration regime, corresponding to the limit (*δ* + *u*)*/b* = *ξ →* 0, as:

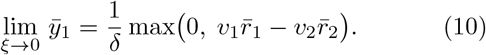

To reconstruct the input-output map (or activation function, used interchangeably throughout this manuscript) for the sequestration-based perceptron, we need to express 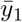 explicitly in terms of the input signals *x*_1_ and *x*_2_, for which we require 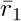 and 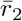. Since a closed-form solution for Eq. (4) is not available, we keep the inputdependent factor *S*_*i*_ from Eq. (6). Using this relation, the fast-sequestration limit becomes

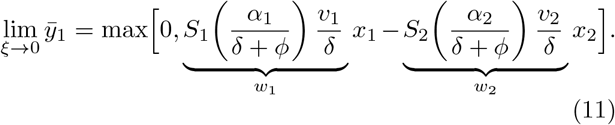

Eq. (11) shows that the sequestration-based perceptron implements a rectified linear unit (ReLU) activation function, in which the inputs *x*_1_ and *x*_2_ are combined through the effective weights *w*_1_ and *w*_2_. Fig. 2-A validates this approximation numerically and shows how the activation function converges to the analytical ReLU prediction from Eq. (11) (white dashed lines) as the sequestration parameter *ξ* decreases. For the rest of the manuscript, we consider a finite *ξ* = 0.01 *µ*M for all numerical simulations, as the thresholding behavior is sufficiently sharp that the fast sequestration approximation is justified [22].

**FIG. 2.**
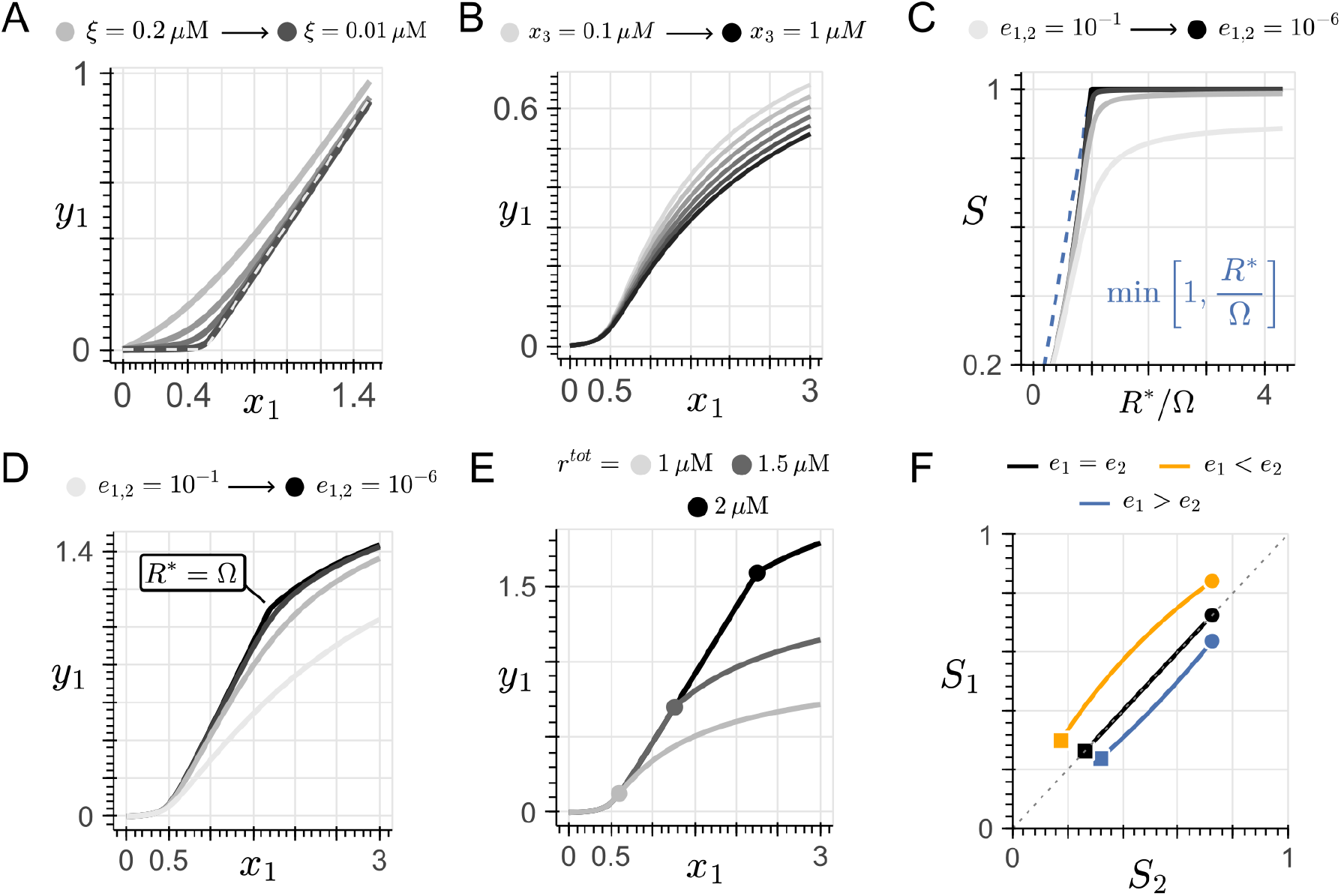
Effect of shared translational resources on the activation function of a sequestration-based perceptron. (A) Numerical validation of the fast-sequestration approximation. The activation function 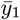 vs *x*_1_ is shown for decreasing values of *ξ* = (*δ* + *u*)*/b*, computed by varying the association rate *b*. As *ξ →* 0, the activation function approximates the analytical ReLU shown in the white dashed line. Darker curves correspond to lower *ξ* values, from 0.2 to 0.01 *µM* . (B) Activation function under increasing endogenous demand, computed by varying *x*_3_ ranging from 0.1 *µM* (lightest) to 1 *µM* (darkest) in the homogeneous case (*e*_1_ = *e*_2_). (C) The ribosome allocation factor *S* as a function of the supply-demand ratio *R*^*\**^*/*Ω for varying mRNA-ribosome binding affinities of the synthetic transcript (light gray to black for *e*_1_ = *e*_2_ ranging from 10^−1^ to 10^−6^). The blue dashed line shows the analytical upper bound min(1, *R*^*\**^*/*Ω) from Eq. (12). (D) Onset of resource competition in the activation function. The strongest binding (darkest) shows a sharp transition at the input value *x*_1_ at which *R*^*\**^ = Ω, while weaker binding (lighter gray) smooths the transition, saturating the output at reduced levels. (E) Location of the onset of resource competition as a function of the total ribosome pool (*r*^*tot*^), ranging from 1 *µM* to 2 *µM* . The circles indicate the onset computed numerically (see Methods). (F) Trajectories in the heterogeneous case of the ribosome allocation factors (*S*_1_, *S*_2_) as *x*_1_ varies from 0 *µM* (squares) to 3 *µM* (circles). The black trajectory along the diagonal represents the homogeneous case (*e*_1_ = *e*_2_), while the blue and orange trajectories show the cases *e*_1_ > *e*_2_ and *e*_1_ < *e*_2_, respectively. Unless stated otherwise, simulations use *x*_2_ = 0.5*µM* and *x*_3_ = 1 *µM* . The total ribosome pool is set to *r*^*tot*^ = 10 *µM* to simulate abundance of resources, to *r*^*tot*^ = 2 *µM* to better display the approximation at higher supply-demand ratios, and swept in Panel E. For the rest of the panels, *r*^*tot*^ = 1 *µM*.

Furthermore, in Eq. (11), each weight is composed of three factors: a transcription term 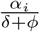, a translation term 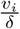, and the ribosome allocation factor *S*_*i*_, for *i ∈* {1, 2}. Here, the transcription and translation terms parametrize protein production for each input, whereas *S*_*i*_ encodes the dependence on the shared translational pool and captures the coupling introduced by resource competition in the activation function. To understand how this dependence influences the behavior of the perceptron, we consider two case studies based on the mRNA–ribosome binding affinities (*e*_*i*_) of the synthetic genes: one in which all mRNAs share the same affinity (the homogeneous case), and another in which the affinities differ (the heterogeneous case). We emphasize that the distinction concerns only the synthetic circuit’s transcripts, since the endogenous demand *M*_*e*_ is treated as a fixed competitor that represents the host organism (see Appendix C).

#### 1. The homogeneous case: identical binding affinities

We define the homogeneous case as one in which all mRNA species from the synthetic circuit bind ribosomes with identical affinity, such that 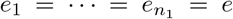. Under this assumption, Eq. (4) can be solved explicitly for the allocation terms *S*_*i*_, which collapse to a single common factor 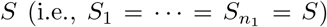. Because all effective weights are multiplied by the same factor *S*, this term can be factored out and acts as an overall multiplicative rescaling of the perceptron weights and, consequently, of the output magnitude. Fig. 2-B illustrates this effect: as the endogenous demand for ribosomes *x*_*e*_ increases (and thus the free available ribosomes decreases), the output 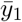 is progressively attenuated (from gray to black in Fig. 2-B; see also Appendix C). However, the thresholding behavior is preserved, and therefore the circuit still behaves as a linear classifier [23].

To understand how the activation function transitions between the resource-abundant regime, in which synthetic circuit can be supplied with enough ribosomes to meet its demand, and the resource-limited regime, in which translational supply becomes a bottleneck, we analyze the closed-form expression for *S* in the strong binding limit (*e →* 0), derived in Appendix D:

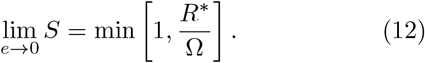

Eq. (12) establishes two regimes based on the ratio *R*^*\**^*/*Ω. When supply exceeds demand (*R*^*\**^ > Ω), *S* = 1 and the output scales linearly with the input, recovering the unattenuated ReLU shown in Fig. 2-A. When demand exceeds supply (*R*^*\**^ < Ω), the output is rescaled by *S* = *R*^*\**^*/*Ω < 1 and saturates as the ribosome pool becomes effectively exhausted. Fig. 2-C plots *S* as a function of the ratio *R*^*\**^*/*Ω for varying binding affinities: as *e →* 0, the curve approximates the upper bound given by Eq. (12) (blue dashed line), while for finite *e* the curves smooth the transition and plateau below *S* = 1. Translating this prediction to the activation function, the strong-binding limit produces a sharp transition in the activation function at the input value *x*_1_ for which *R*^*\**^ = Ω, beyond which the output transitions from linear into the resource-limited regime, where it saturates as 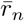 depletes (darkest curve in Fig. 2-D). We refer to this transition as the onset of resource competition. On the other hand, for finite binding affinity (lighter gray curves in Fig. 2-D), weaker ribosome binding prevents full utilization of the available pool, leading to *S* < 1 even when *R*^*\**^ > Ω. As a result, the sharp transition between regimes is smoothed, and the output exhibits gradual saturation at reduced levels despite abundant ribosomes.

Having defined the onset of resource competition, we now ask what determines its location along the activation function. We restrict our analysis to the regime in which the synthetic transcripts bind to ribosomes more strongly than the endogenous one (*e*_1_, *e*_2_ *≪ e*_3_), as this is a design regime accessible through ribosome binding site (RBS) engineering, and the one in which the circuit is assumed to operate. In this regime, the input value *x*_1_ at which the onset occurs depends on the total ribosome pool (*r*^*tot*^) and on the synthetic circuit transcription parameters, and admits a closed-form solution (see Appendix E). Fig. 2-E illustrates this effect: as *r*^*tot*^ decreases, the onset shifts toward smaller input values *x*_1_. In Appendix E, we further characterize how the onset shifts when the binding-affinity ordering between the synthetic and endogenous transcripts is relaxed, and found that in regimes where the endogenous transcript binds comparably or more strongly than the synthetic ones to ribosomes, the location of the onset becomes sensitive to *x*_*e*_ and *e*_3_ as well.

### 2. The heterogeneous case: varied binding affinities

In experimental implementations, it is challenging to ensure homogeneous mRNA–ribosome affinities across all components of a synthetic circuit, and some degree of heterogeneity is typically present [24]. Although Eq. (4) cannot be solved analytically in this regime, we can analyze how the rescaling terms *S*_1_ and *S*_2_ appear in Eq. (11) and shape the resulting activation function. Fig. 2-F plots the parametric trajectory of (*S*_1_, *S*_2_) as the input *x*_1_ varies. In the homogeneous case (*e*_1_ = *e*_2_, black line), both transcripts exert identical demand on the ribosome pool, producing a trajectory that traces the diagonal *S*_1_ = *S*_2_ across all levels of ribosome availability. When *M*_1_ has weaker ribosome affinity (*e*_1_ > *e*_2_, blue line), its translation is more strongly attenuated as free ribosomes are depleted. As competition increases, *S*_1_ decreases more rapidly than *S*_2_, causing the trajectory to fall below the diagonal and indicating that *M*_1_ receives a smaller allocation of the available ribosomes than *M*_2_ in the resource-limited regime. Conversely, when *M*_1_ binds ribosomes more strongly (*e*_1_ < *e*_2_, orange line), it maintains larger *S*_1_ values under the same conditions, shifting the curve above the diagonal and reflecting a larger share of the ribosome pool under competition.

Notably, despite differential rescaling by the *S*_*i*_ factors in the heterogeneous case, the thresholding behavior of Eq. (11) is preserved across all binding-affinity scenarios. Resource competition modifies the relative magnitudes of the two effective weights but does not eliminate the nonlinearity itself. Because this thresholding behavior is required for classification [23], our analysis indicates that, under heterogeneous binding affinities, the sequestration-based perceptron continues to operate as such.

### C. Quantifying the effect of shared translational resources on the decision boundary

The rescaling of the effective weights due to ribosome competition shifts the location and orientation of the decision boundary. Here, we analyze these shifts and characterize the resulting changes in its geometry.

#### 1. Molecular perceptron under shared translational resources

We can write an explicit expression for the decision boundary of a single perceptron as

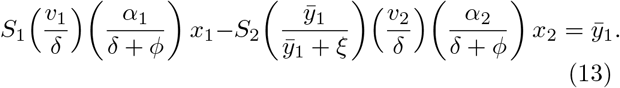

This expression defines a linear decision boundary in the input space (i.e., (*x*_1_, *x*_2_) plane), whose geometry is shaped by the ribosome allocation terms *S*_1_ and *S*_2_. To characterize this geometry, we define the slope of the decision boundary (*Q*) as

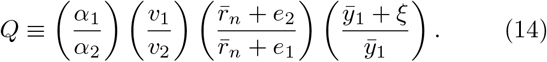

This expression can be further decomposed into the following four terms: (I) the ratio of transcription rates (*α*_1_*/α*_2_); (II) the ratio of translation rates (*v*_1_*/v*_2_); (III) the relative efficiency of ribosome utilization, 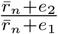 (or, equivalently, 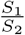); and (IV) the dissociation constant *ξ*. In the ideal case where the two inputs have identical transcription and translation rates, ribosomes are abundant 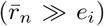, and sequestration is fast (*ξ →* 0), we obtain *Q →* 1, corresponding to a diagonal decision boundary, as shown in Fig. 3-A. Under moderate and extreme ribosomal competition (i.e., increased endogenous demand) in the homogeneous-affinity case, the decision boundary remains unchanged because the slope *Q* is preserved when *e*_1_ = *e*_2_, although the output amplitude 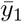 is reduced, as shown in Fig. 3-B and C, respectively.

**FIG. 3.**
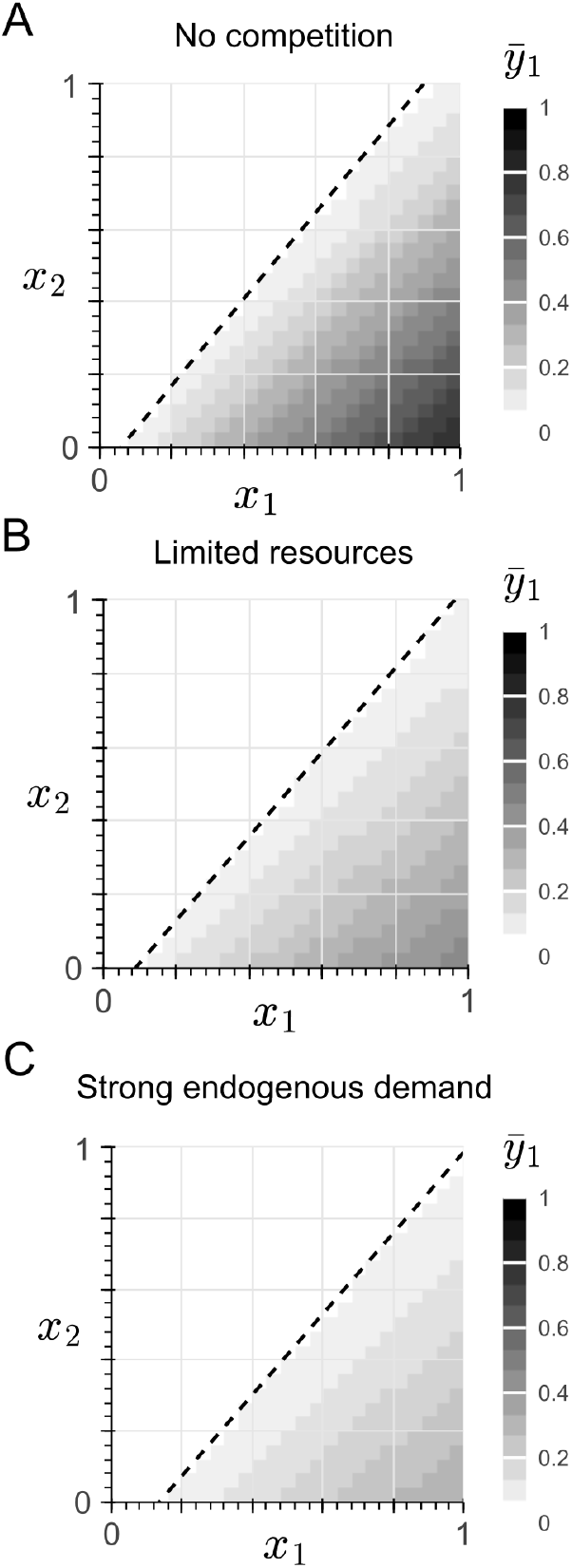
Effect of shared translational resources on the decision boundary of a sequestration-based perceptron under equal mRNA-ribosome binding affinities (homogeneous). For a total ribosome pool of *r*^tot^ = 1.8 *µ*M, the heatmaps in the input space show the steady-state output 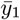 across three resource-availability regimes: (A) no resource competition (*x*_*e*_ = 0 *µ*M), (B) limited competition in the homogeneous-affinity case (*x*_*e*_ = 3 *µ*M), and (C) high endogenous demand (*x*_*e*_ = 5 *µ*M).

Furthermore, to analyze the heterogeneous-affinity case, we define the ratio between the rescaling terms as

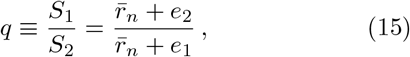

Then, we can rewrite Eq. (13) as follows:

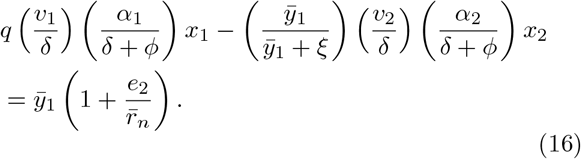

From this expression, we note that the effect of heterogeneous ribosome affinities is captured by the factor *q*, which modifies the slope of the linear decision boundary, and from which we can identify three regimes. In the first regime, translational resources are abundant 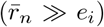, leading to *q →* 1, such that affinity heterogeneity does not affect the decision boundary. In the second regime, where 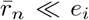, the ratio approaches *q → e*_2_*/e*_1_. However, achieving 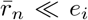 would require either unrealistically large inputs or transcriptional weights, or an extremely small total ribosome supply. We therefore focus our analysis on the third regime, where 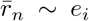, and affinity heterogeneity induces changes in the slope of the decision boundary.

In this regime, given that *q* is implicitly dependent on the inputs *x*_1_ and *x*_2_, Eq. (16) describes a nonlinear decision boundary. To determine the orientation of the boundary’s curvature, we compute the partial derivatives of *q* with respect to each input *x*_*i*_, treating 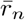 as an implicit function of 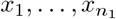 defined by Eq. (4). This yields

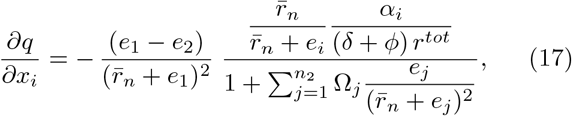

for *i* = 1, … , *n*_1_. Because all remaining terms in Eq. (17) are strictly positive, the sign of *∂q/∂x*_*i*_ is determined solely by the difference *e*_1_ − *e*_2_. When *e*_1_ < *e*_2_, the derivative is positive and *q* increases with increasing inputs, resulting in an upward curvature of the decision boundary. Conversely, when *e*_1_ > *e*_2_, the derivative is negative and *q* decreases as the inputs increase, producing a downward curvature.

To illustrate this behavior, we computed the decision boundary for a representative set of parameters: *r*^tot^ = 1.8 *µ*M and an endogenous demand *x*_3_ = 1 *µ*M. These parameters were chosen to make the curvature of the decision boundary easy to identify within the input space, yet the qualitative results detailed next are independent of this choice, as shown in Appendix F. Moreover, we consider that the endogenous transcript *M*_*e*_ binds ribosomes with affinity comparable to the synthetic ones (*e*_3_ *∼ e*_1,2_). In this regime, the endogenous transcript depletes the ribosome pool sufficiently for the decision boundary to be in the regime 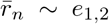 and enables the predictions from Eq. (17). This behavior is illustrated in Fig. 4-A1 and A2, which show the upward and downward bending. The decision boundary remains approximately linear while free ribosomes are abundant, and begins to bend as the system enters the regime in which ribosomes become limiting. This change in geometry coincides with the system crossing the onset of resource competition. Fig. 4-B shows the free ribosome fraction 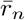 as a heatmap, with the locus *R*^*\**^ = Ω overlaid. The heatmap transitions from a blue gradient (abundant free ribosomes) to white (depleted ribosome pool) as the system crosses this locus, whose location is determined by the total ribosome pool. From Eq. (5), varying *r*^*tot*^ shifts the onset towards the origin for smaller values, and away from the origin for larger ones (see Appendix F for further details).

**FIG. 4.**
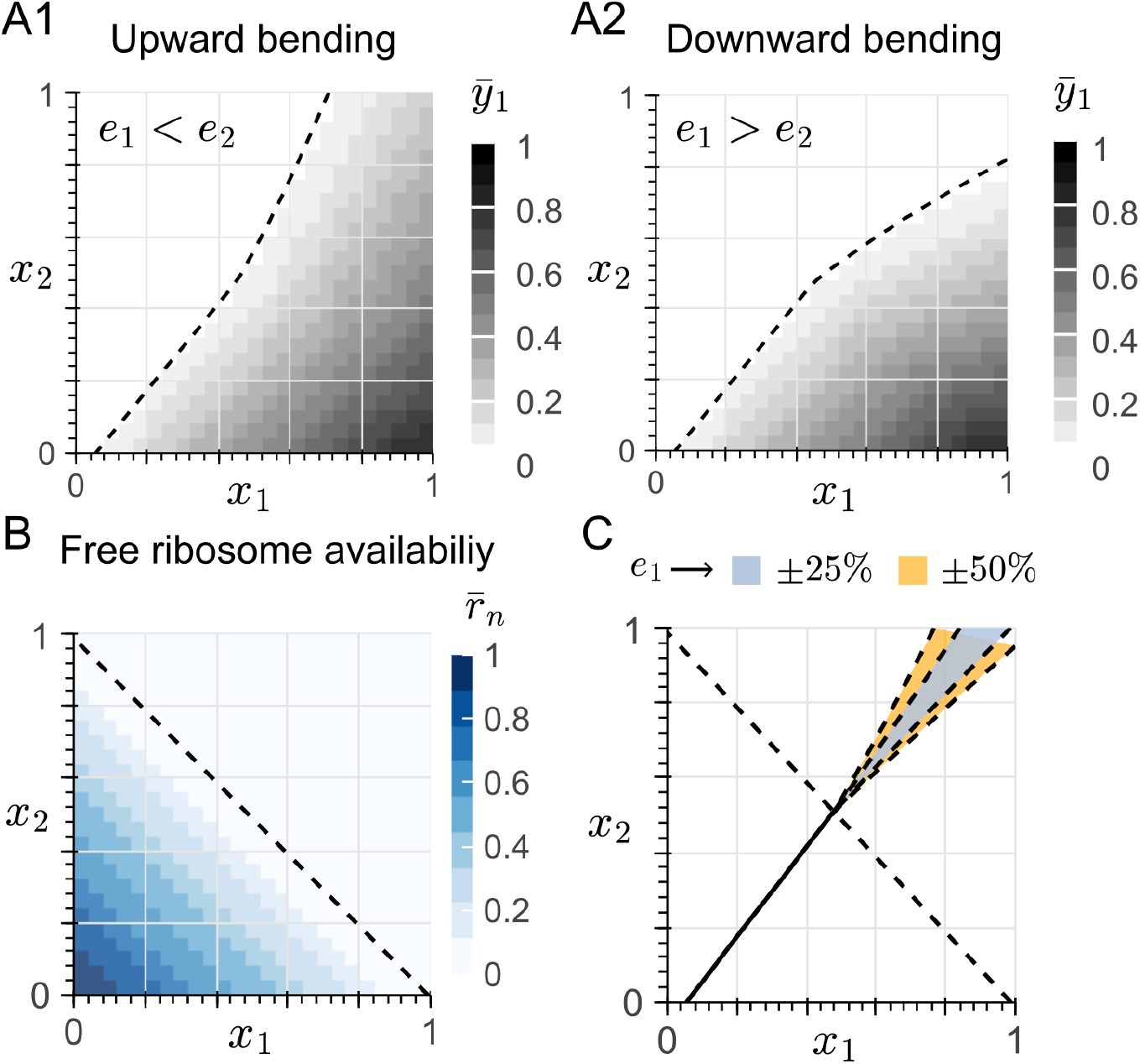
Effect of shared translational resources on the decision boundary of a sequestration-based perceptron under different mRNA-ribosome binding affinities (heterogeneous). (A1) Steady-state output 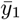 in the input space for the heterogeneous case *e*_1_ < *e*_2_ (i.e., *M*_1_ binds ribosomes more strongly than *M*_2_). The black dashed line shows the onset locus of resource competition *R*^*\**^ = Ω. (A2) Same as Panel A1, for the opposite heterogeneous case *e*_1_ > *e*_2_. (B) Free ribosome fraction 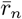 across the input space. The black dashed line is the onset locus *R*^*\**^ = Ω. (C) Uncertainty region of the decision boundary under a bounded variation in the ribosome-binding affinities. The blue shaded region corresponds to the envelope for *e*_1_ *±* 25% variations of the value used in Panel A1, whereas the orange shaded region corresponds to the envelope for a higher *±*50% variation. All panels use *r* = 1.8 *µM* , *x*_3_ = 1 *µM* , and a baseline affinity *e*_*i*_ *≈* 6.5 × 10.

If we then vary the ribosome-binding affinities within a bounded interval, each parameter choice generates a slightly different decision boundary. The collection of these boundaries forms an envelope in the input space, which we define as an uncertainty region (Fig. 4-C). For variations of *±*25% and *±*50% around a baseline binding affinity value *e*_*i*_, the envelope is contained within a predictable region of the input space, with its width set by the magnitude of the affinity variation. Because binding affinities cannot be tuned exactly in practice, the realized boundary may lie anywhere within this envelope, rendering points inside it ambiguous. Importantly, the envelope itself remains bounded by the predefined variability in *e*_*i*_, so even under strong resource limitation the boundary stays confined within this region.

#### 2. Molecular multi-layer perceptron (MLP) under shared translational resources

Building on the analysis of the single perceptron, we study how shared translational resources affect the decision boundary of a multi-layer perceptron (MLP). As a representative case, we consider an MLP with two nodes in the hidden layer and a single output node, with weights and biases chosen to generate distinct decision boundaries. Specifically, Fig. 5-A1 implements a closed, triangular-like decision boundary. In this architecture, the output of the final node is the protein complex formed through molecular sequestration, following a previously proposed design [25]. By contrast, Fig. 5-B1 generates a non-convex decision boundary, where the output of the final node is the free (unbound) protein. The full dynamical models for both architectures are provided in Appendix G.

**FIG. 5.**
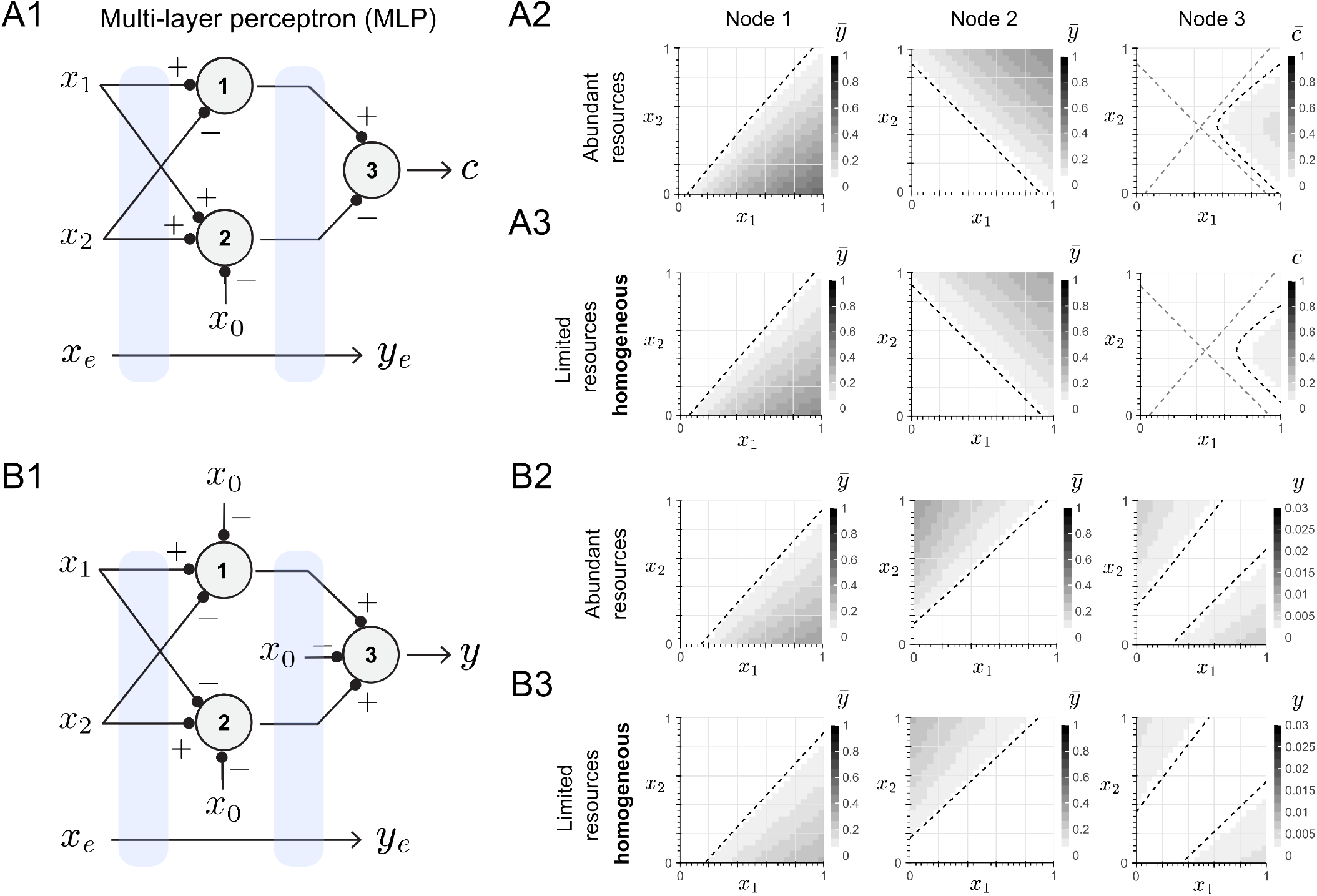
Effect of shared translational resources on the decision boundary of a multi-layer perceptron (MLP) in the homogeneous-affinity regime. (A1) and (B1) show an MLP architecture with two nodes in the hidden layer and one output node, with weights chosen to produce different decision boundaries (A: triangular pattern, B: non-convex pattern). At each node, two proteins are produced and interact via molecular sequestration. For Nodes 1 and 2, the output is the free protein, as in the single-perceptron case. For Node 3, the output is the protein complex in (A1), whereas in (B1) the output is again the free protein. All mRNA species compete for a shared ribosome pool of *r*^tot^ = 3 *µ*M in (A) and *r*^*tot*^ = 2 *µ*M in , as indicated by the blue bands on the weights. (A2) and (B2) correspond to the abundant-resource regime (*x*_*e*_ = 0 *µ*M), where resource competition is negligible. (A3) and (B3) show the limited-resource, homogeneous-affinity case (*x*_*e*_ = 1.5 *µ*M), in which all mRNA species have identical ribosome-binding affinities. For (A2) and (A3), the faint dashed lines represent the linear decision boundaries of Node 1 and Node 2 overlapped in the input space in Node 3. For (B2) and (B3), the decision boundary of Node 3 exactly overlaps those from the previous nodes.

Before analyzing the MLP decision boundary under translational resource constraints, we extend the singleperceptron formulation to include a bias species, as illustrated in Node 2 of Fig. 5-A1. In this architecture, the inputs *X*_1_ and *X*_2_ each produce an mRNA molecule (*M*_1_ and *M*_2_) that translate into the same protein *Y* , whereas the bias input *X*_0_ produces a third mRNA *M*_0_ that translates into the protein responsible for sequestering *Y* . If the ribosome-binding affinity of *M*_0_ changes relative to *M*_1_ and *M*_2_, the bias term effectively sets a fixed threshold in the boundary equation (see Appendix H). Changes in *M*_0_ therefore translate the decision boundary without bending it. By contrast, if *M*_1_ and *M*_2_ have different ribosome-binding affinities – for example due to distinct RBS even though they encode the same protein –, the relative contribution of the two inputs could, in principle, change the slope of the boundary. However, because both inputs produce the same protein, movement along the boundary requires that an increase in one input be compensated by a decrease in the other. This compensation keeps the total ribosome demand from *M*_1_ and *M*_2_ approximately constant for the threshold set by *M*_0_. As a result, free ribosome levels vary little along the boundary, and it appears as a tilted straight line rather than a curved one.

Returning to MLPs, unlike the single perceptron case, it is challenging to obtain a closed-form expression for the nonlinear decision boundary generated by the network. We therefore analyze it numerically. Fig. 5-A2 and B2 show the expected decision boundaries for both architectures under abundant translational resources, where ribosome competition is negligible across all nodes of the MLP. Under these conditions, each node behaves approximately as an isolated perceptron, and the output decision boundary can be interpreted as a nonlinear composition of the linear boundaries generated by the intermediate nodes.

Note that the output magnitude of the MLP in Fig. 5-B1 is smaller than in Fig. 5-A1 for two reasons. First, the total number of ribosomes engaged in translation differs between the architectures (8 complexes in Fig. 5-A1 versus 10 in Fig. 5-B1; see Appendix G). Second, the output variables are defined differently. In Fig. 5-A1, the output *c* takes the minimum value between the total concentrations of its two inputs [22]. In contrast, in Fig. 5-B1 the output *y* computes the difference between inputs assigned positive weights and those assigned negative weights, and then applies a max operator [21]. This operation intrinsically produces a smaller output magnitude.

We next examine the effect of ribosome competition on the decision boundaries of the MLP. For the homogeneous-affinity case, Fig. 5-A3 and B3 show that limited resources uniformly rescale the output magnitude while preserving the overall shape of the decision boundaries across all three nodes. In contrast, Fig. 6-A1 and B1 show that heterogeneity in the mRNA-ribosome binding affinity bends the decision boundaries of Node 1 and Node 2, and these distortions propagate to the output node. We also computed uncertainty regions to indicate where the decision boundary shifts when the binding affinities *e*_*i*_ vary within a *±*50% interval around their nominal values. Fig. 6-A2 and B2 illustrate how these uncertainties propagate with network depth. The upper half of Node 3’s uncertainty region originates from Node 1 (blue), while the lower half originates from Node 2 (orange). As in the single-perceptron case, for a predefined variability in *e*_*i*_ the envelope of the uncertainty region remains bounded and therefore predictable.

**FIG. 6.**
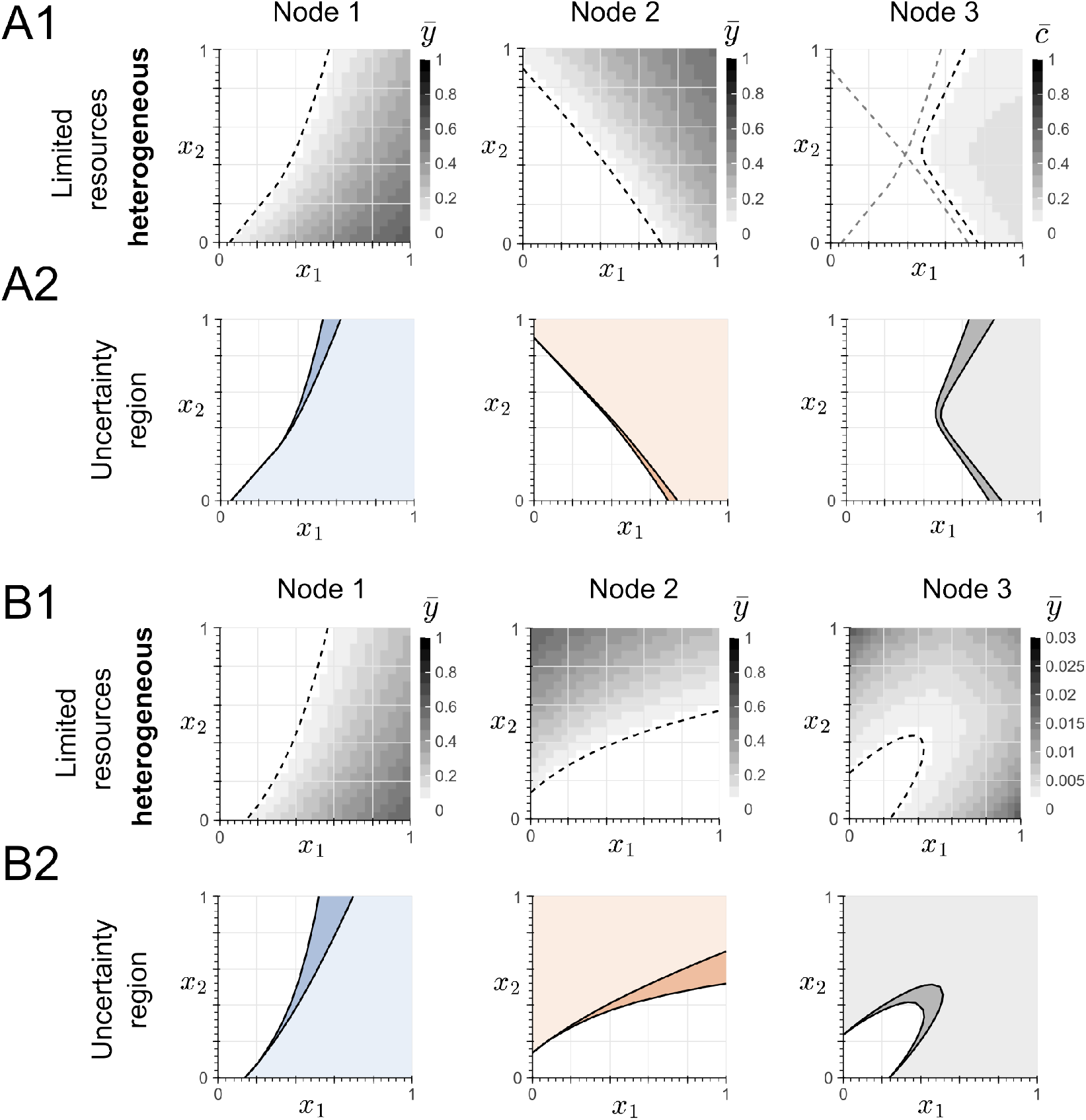
Effect of shared translational resources on the decision boundary of a multi-layer perceptron (MLP) in the heterogeneous-affinity regime. (A1) and (B1) show the limited-resource, heterogeneous-affinity case for a shared ribosome pool of *r*^tot^ = 3 *µ*M, with nominal ribosome-binding affinity *e*_*i*_ = 6.5 × 10^−6^. In both cases, the affinity of the mRNA produced by *X*_1_ (i.e., *e*_1_) is set to 0.2 times this nominal value. (A2) and (B2) show the corresponding uncertainty regions for each node. Light shading indicates outputs above threshold in both endpoint simulations, darker shading indicates disagreement between the two simulations, and white indicates outputs below threshold in both. Solid black curves mark the threshold contours. The simulations in (A) and (B) correspond to the MLP architectures shown in Fig. 5.

### D. Extending the analysis to growth-mediated feedback

Throughout this work, we have treated the total ribo-some pool *r*^*tot*^ as a fixed parameter, yet in practice, its value is coupled to the physiology of the cell. The expression of a synthetic gene such as the sequestration-based perceptron diverts resources from essential cellular processes, imposing burden on the cell that is observed as a reduction in growth. Consequently, the global translational capacity and dilution rate of every chemical species change accordingly [14]. To introduce growth feedback into our framework, we included two minimal modifications without committing to a coarse-grained proteome allocation model [19, 26] or to an empirical growth law [12]. First, we replace the degradation rate *δ* with the growth rate *λ* in Eqs. (1)–(3), since dilution is a consequence of cell division. Second, we treat *r*^*tot*^ as a state variable rather than a fixed constant. At steady state, both quantities take the values *λ*^*\**^ and *r*^*tot,\**^, determined by the total translational burden Ψ, so that 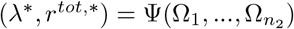. We do not specify the functional form of Ψ as it is outside the scope of this work, and refer the reader to the following literature for further details: [12, 19, 27, 28].

Under these modifications, the steady state derivations detailed in Appendix A and B carry through, and the dimensionless parameters *e*_*i*_ and Ω_*i*_ acquire an implicit dependency on *λ*^*\**^ and *r*^*tot,\**^. We refer to a derivation as form-invariant under growth feedback if it retains the same algebraic relations between the quantities 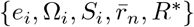. Regarding the activation function, replacing *δ → λ* and *r*^*tot*^ *→ r*^*tot,\**^ in Eq. (11) results in

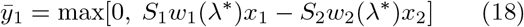

The effective weights *w*_*i*_(*λ*^*\**^) = *α*_*i*_*v*_*i*_*/*[*λ*^*\**^(*λ*^*\**^ + *ϕ*)] now depend on the steady-state growth rate, and the ribosome allocation factors *S*_*i*_ are a function of (*λ*^*\**^, *r*^*tot,\**^). Growth-mediated feedback scales the output 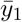 and shifts the location of the threshold, but the thresholding behavior of Eq. (18) is preserved. Moreover, the equation for the decision boundary becomes

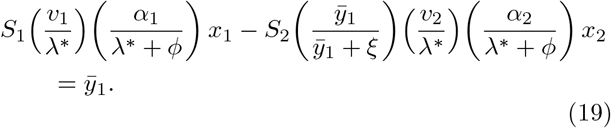

Eq. (19) has the same structure as Eq. (13), with *δ* replaced by *λ*^*\**^ and the ribosome allocation factors *S*_*i*_ now evaluated at the input-dependent values of *λ*^*\**^ and *r*^*tot,\**^.

In the homogeneous case, for *e*_2_ = *e*_1_, both transcripts share the ribosome allocation factor *S* = *S*(*λ*^*\**^, *r*^*tot,\**^), and the slope of the decision boundary reduces to *Q* = *α*_1_*v*_1_*/*(*α*_2_*v*_2_), independent of *λ*^*\**^ and *r*^*tot,\**^. The geometry of the decision boundary is therefore unchanged in this regime. The effect of growth can be observed in the effective weights *w*_*i*_, which now include the term *α*_*i*_*v*_*i*_*/*[*λ*^*\**^(*λ*^*\**^ + *ϕ*)] multiplied by *S*. This solely attenuates the output by a factor that depends on *λ*^*\**^ (through *w*_*i*_) and on *r*^*tot,\**^ (through *S* via *e*_*i*_ and 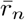).

For the heterogeneous case (*e*_2_ ≠ *e*_1_), even though the structural form of Eq. (19) is preserved, growth-mediated feedback results in the bending of the decision boundary with a direction no longer predictable from the affinity ordering alone. As detailed in Appendix I, the direction depends on how the host’s physiology couples translational burden to the ribosome pool. However, since *λ*^*\**^ and *r*^tot,*\**^ take values within a physiologically bounded range for any host [29], the resulting uncertainty in the decision boundary remains bounded to a region of the input space.

Furthermore, in this same regime and for a sequestration-based perceptron with two inputs, we can identify the transition from the resource-abundant to the resource-limited regime by the condition Ω = *R*^*\**^, which becomes the following implicit equation under growth feedback:

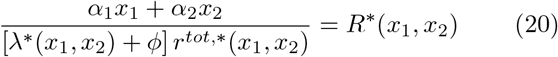

When *r*^*tot,\**^ is treated as a constant, as in Section II.B.2, the denominator of Eq. (20) is fixed and the onset locus of resource competition is a straight line in the input space, illustrated in Fig. 4-B. Under growth feedback, whether this locus bends or remains straight depends on the structure of Ψ. For growth laws in which Ψ depends only on the total burden Σ_*i*_ Ω_*i*_ (e.g., [14]), the onset condition itself fixes the value of total burden, and therefore the values of *λ*^*\**^ and *r*^*tot,\**^ along the locus. In other words, the onset locus remains a straight line. For more general Ψ that details the burden between individual transcripts (e.g., [19]), *λ*^*\**^ and *r*^*tot,\**^ vary along the locus, which bends and causes the onset for lower input magnitudes. Further details are included in Appendix I.

## III. DISCUSSION

In this work, we developed a resource-aware framework to analyze how competition for translational resources affects the decision boundaries of sequestration-based neural networks operating at the protein level. Our analysis shows that, in the homogeneous-affinity regime, ribosome competition rescales the output magnitude without altering the geometry of the decision boundary, preserving the classifier’s function. In contrast, when mRNA-ribosome binding affinities are heterogeneous, differential affinities introduce curvature in the decision boundary, whose direction is determined by the sign of the affinity difference. Importantly, for a predefined variability in binding affinities, the resulting bending of the decision boundary remains confined within a bounded uncertainty region. As a result, the possible decision boundaries remain constrained, which suggests that the classifier’s behavior may remain predictable. These results hold for both single perceptrons and multilayer architectures, where distortions generated at hidden nodes propagate through the network and combine at the output layer. Although our two-layer results show that this propagation remains limited, deeper networks may accumulate larger uncertainty regions as distortions from successive layers compound. Understanding how these effects scale with network depth remains an important direction for future work.

Although our analysis focused on molecular sequestration reactions, previous studies show that synthetic circuits based on catalytic degradation [25], phosphorylation/dephosphorylation [30] and competitive binding [23] can also operate as perceptrons. We hypothesize that our conclusions extend to these architectures as well, because translational resource competition primarily applies a multiplicative rescaling to the effective weights through the ribosome allocation factor *S*_*i*_. In the homogeneous regime, this rescaling uniformly attenuates all weights, preserving the decision boundary geometry while reducing the output magnitude. Likewise, in the heterogeneous case, a predefined variability in the mRNA-ribosome affinity results in a bounded envelope for the uncertainty region regardless of the specific non-linearity used for thresholding.

From a design perspective, these observations suggest practical guidelines for implementing molecular classifiers at the protein level. While the total pool of ribo-somes is largely determined by the host [29, 31], competition effects can be mitigated by strategies that reduce demand. For example, prior work has developed feed-back controllers, including negative feedback and feedforward loop strategies, that adjust mRNA expression to maintain ribosome demand below limiting levels [32–40]. Here, Eq. (4) indicates that weakening ribosome binding reduces sequestration of free ribosomes and thereby lowers demand, a strategy consistent with more detailed models of translation [13, 41]. In practice, this can be achieved by using a weaker RBS or otherwise lowering translation-initiation rates. Another complementary approach is to employ orthogonal ribosome systems, which provide a dedicated translational pool for the synthetic circuit and reduce coupling to host gene expression [42, 43]. This partially decouples the ribosomes used for endogenous transcripts from those available to the circuit (i.e., a design regime around Eq. (4) for *R*^*\**^ = 1). In this scheme, if all circuit transcripts could achieve the same ribosome-binding affinity, the effect on output magnitude and the decision boundary would be minimal. However, in practice, identical affinities are difficult to achieve: even with the same RBS, effective binding depends on the transcript’s secondary structure and local sequence context [44]. Nevertheless, our analysis shows that the resulting deviations remain bounded, suggesting that robust classifier design is feasible despite this unavoidable heterogeneity.

In this study, we considered that each input produces a distinct mRNA molecule, a design choice that allows us to explicitly study the effect of multiple transcripts competing for the shared ribosome pool. This formulation captures a general scenario in which every synthetic gene independently draws on the translational machinery. Interestingly, prokaryotic cells face an analogous situation: endogenous mRNAs possess diverse ribosome binding site strengths, and therefore exhibit heterogeneous ribosome affinities [44, 45]. Our analysis shows that such heterogeneity can generate nonlinear decision boundaries as a direct consequence of the coupling introduced by resource competition, without requiring a separate regulatory mechanism. This observation leads to questions such as whether cells exploit this effect for information processing using fewer computational parts, or if they employ mitigation strategies instead. Cell-free systems offer a particularly interesting experimental platform for rapidly validating these predictions, as they allow direct control over ribosome concentrations, systematic variation of RBS strengths, and the possibility of decoupling transcription from translation by independently adjusting RNA polymerase and ribosome levels [46].

Future work will extend this framework along several directions. A more comprehensive model of the cellular environment would incorporate competition for RNA polymerase, growth-mediated feedback on global transcriptional and translational capacity, and shared RNA degradation pathways. In particular, bacterial growth laws establish a direct link between ribosome allocation and growth rate [14]. While Section II.D shows that the structural conclusions of our framework are preserved under a phenomenological treatment of growth feedback, incorporating a specific coarse-grained proteome allocation model [19, 26] or empirical growth law [12] would allow quantitative predictions of how the location and curvature of the decision boundary shift with the host’s physiological state. Finally, scaling the analysis to larger networks with additional hidden layers would help to understand how uncertainty regions propagate with network depth and inform the maximum expressivity of molecular classifiers under shared translational resources.

## IV. METHODOLOGY

### A. Numerical integration

All simulations were performed in Python using scipy.integrate.odeint. Each system of ODEs was integrated from zero initial conditions for all species over a time window of *t* = 200 hours, and the final value of the numerical solution was taken as the steady-state concentration. To construct the activation function of a single perceptron, one input was held fixed (e.g., *x*_2_ = 0.5 *µ*M) while the other was varied over a specified range, and the steady-state output concentration was recorded for each value. To construct the heatmaps in the (*x*_1_, *x*_2_) plane, we generated a 25 × 25 grid of input combinations, with each input varying from 0 to 1 *µ*M. The decision boundary was extracted from these heatmaps as the contour line at a threshold value of 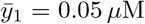, corresponding to the first visible transition in the output (i.e., white to light color in the heatmaps). This threshold was chosen because, for a finite sequestration rate, the steady-state output at the perceptron’s nominal threshold (i.e., where 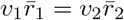) is not zero but rather

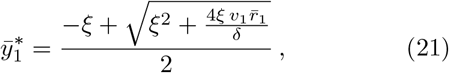

which depends on the ribosome complex concentration 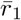 and thus varies across the input space. A fixed numerical threshold was therefore used in place of the inputdependent analytical value. Contour extraction was performed using matplotlib.pyplot.contour.

### B. Uncertainty regions

The uncertainty regions shown throughout the manuscript were computed by varying the mRNA-ribosome binding affinities *e*_*i*_ within a specified percentage interval around their nominal values. For each sampled set of affinities, the steady-state output was computed over the full input grid and the corresponding decision boundary was extracted. The envelope of all such boundaries defines the uncertainty region in the input space.

### C. Model parameters

Parameters were chosen to reflect representative values for synthetic gene expression in *E. coli*, obtained or inferred from [13, 47]. Transcription rate constants of *α*_*i*_ = 10 hr^−1^ give an mRNA production rate of 10 *µ*M hr^−1^ at an input concentration of *x*_*i*_ = 1 *µ*M, corresponding to approximately 6,000 transcripts per cell per hour (using the conversion 1 *µ*M *≈* 600 molecules per *E. coli* cell), consistent with a strong promoter on a medium-to-high copy number plasmid (approximately 100 transcripts per plasmid per hour for a *∼* 60-copy plasmid).

The mRNA degradation rate *ϕ* = 10 hr^−1^ gives a halflife of approximately 4.2 minutes, typical for *E. coli* transcripts, while the dilution rate *δ* = 1 hr^−1^ corresponds to a doubling time of approximately 42 minutes. The ribosome–mRNA association and dissociation rates (*a*_*i*_ = 100 *µ*M^−1^ hr^−1^, *d*_*i*_ = 1 hr^−1^) yield an effective dissociation constant *K*_*d*_ = *d*_*i*_*/a*_*i*_ = 0.01 *µ*M = 10 nM, representative of a strong ribosome binding site.

Translation rates of *v*_*i*_ = 1 hr^−1^ represent the effective rate of protein production per ribosome–mRNA complex, incorporating initiation inefficiency and elongation pauses. The sequestration parameters (*b* = 200 *µ*M^−1^ hr^−1^, *u* = 1 hr^−1^) give *K*_*d*_ = *u/b* = 0.005 *µ*M = 5 nM, consistent with strong protein–protein binding. Unless otherwise specified in the figure captions, the total ribosome pool *r*^tot^ represents the fraction of ribosomes available to the synthetic circuit, set to 1 *µ*M for the single perceptron and 3 *µ*M for the MLP architectures to account for the larger number of transcripts competing for resources.

## ACKNOWLEDGMENTS

We thank Kirill Sechkar and Nicholas Nolan for their constructive comments on the manuscript, and the members of the Cuba Samaniego Laboratory for their insights during the conceptualization of this work.

## DATA AVAILABILITY

The code that supports the simulations of this manuscript is openly available in the GitHub repository.

## Appendix A: Steady-state analysis of protein production under ribosome competition

We begin by stating the mass conservation relation for the total ribosome concentration, which enforces that the concentration of free ribosomes (*R*) together with the concentrations of ribosomes bound in translation complexes (*R*_*i*_) remains constant. We explicitly distinguish between ribosomes bound to synthetic mRNAs and those bound to endogenous transcripts, resulting in the following equation:

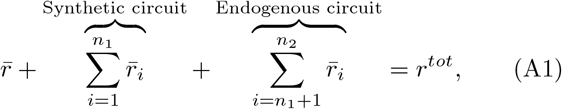

where 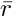 and 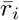 denote the steady-state values of *r*(*t*) and *r*_*i*_(*t*), respectively. By setting the right-hand sides of Eqs. (1) and (2) to zero, we solve for the steady-state mRNA concentration 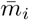 as a function of 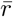 and 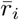:

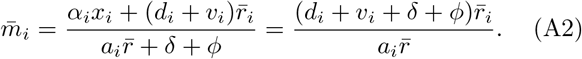

This expression allows us to solve for the steady-state ribosome–mRNA complex, resulting in the following equation:

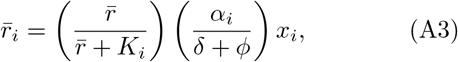

where

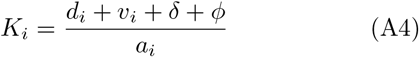

represents the dissociation constant between *M*_*i*_ and the ribosome. Moreover, for convenience, we normalize by the total ribosome pool *r*^*tot*^ and define the following dimensionless parameters:

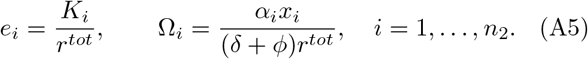

By introducing 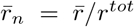 to denote the normalized steady-state concentration of free ribosomes, we can express the ribosome conservation equation in a simplified form as written in Eq. (4), obtained by dividing Eq. (A3) by *r*^*tot*^, replacing with the dimensionless variables from Eq. (A5), and replacing in Eq. (A1).

## Appendix B: Steady-state analysis of sequestration-based perceptron

By setting Eq. (9) to zero and solving the steady-state of Eqs. (7) and (8) in terms of 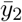, we obtain

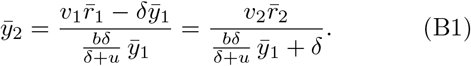

where 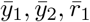 and 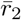 are the steady-state values of *y*_1_(*t*), *y*_2_(*t*), *r*_1_(*t*) and *r*_2_(*t*), respectively. Expanding Eq. (B1), we can write a second order polynomial in terms of 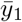 of the shape 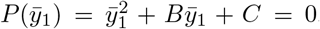. Let *ξ* = (*δ* + *u*)*/b*, so 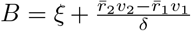 and 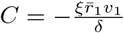. The quadratic admits a single solution due to the non negativeness of chemical concentrations, given by:

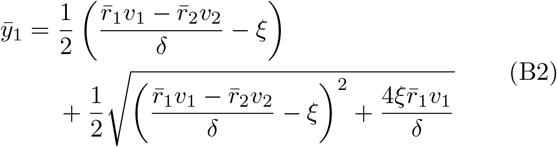

Following the fast-sequestration approximation used in [21, 22], in the regime 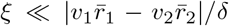, Eq. (B2) follows the reduced expression obtained in the fast-sequestration limit written in Eq. (10). On the other hand, the equation for the decision boundary shown in Eq. (13) can be obtained from Eq. (B1) by assuming 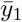 as a constant, and writing 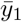 as a function of the inputs *x*_1_ and *x*_2_

## Appendix C: Phenomenological modeling of the endogenous demand

Throughout this manuscript, we model the endogenous translational demand as a single mRNA species *M*_*e*_ that is produced at a level determined by the input *x*_3_ and with ribosome-binding affinity *e*_3_. This is the minimal phenomenological model chosen so that *x*_3_ and *e*_3_ can be considered summary statistic of the several mRNAs that simultaneously compete for the ribosome pool in bacteria, each with their own copy number, transcription and translational rates, and ribosome-binding affinities [48]. In this way, *x*_3_ and *e*_3_ lump the aggregate demand of all such individual mRNAs into the dimensionless quantity *S*_3_Ω_3_ that is included in the ribosome conservation equation (Eq. (4)) through the definition of *R*^*\**^ in Eq. (5).

In this way, we can study how the endogenous demand affects the synthetic circuit’s behavior as a classifier in terms of *x*_3_ and *e*_3_. In particular, Fig. 2-B shows how increasing *x*_3_ attenuates the perceptron’s output while preserving the thresholding behavior. The same qualitative effect can be obtained by increasing the endogenous transcript’s binding affinity, or, more globally, by reducing the total ribosome pool. Both alternatives lower the supply-demand ratio *R*^*\**^*/*Ω that determines the magnitude of the ribosome allocation factor in Eq. (12). In Fig. 7-A, for a fixed *x*_3_ input value, we sweep *e*_3_ across values starting from the same affinity as the synthetic transcript (*e*_3_ = 10^−6^ = *e*_1_ = *e*_2_) to a thousand-fold increase in affinity (for visualization purposes, *e*_3_ = 10^−9^), showing how the perceptron’s output gets attenuated across the entire input range. Fig. 7-B, in turn, sweeps the total ribosome pool *r*^*tot*^ for a fixed endogenous demand (*x*_3_ = 1 *µM* , *e*_3_ = 100), showing that reducing the supply of ribosomes produces the same qualitative attenuation. Importantly, in all cases, the perceptron’s attenuation in magnitude does not alter the thresholding behavior.

**FIG. 7.**
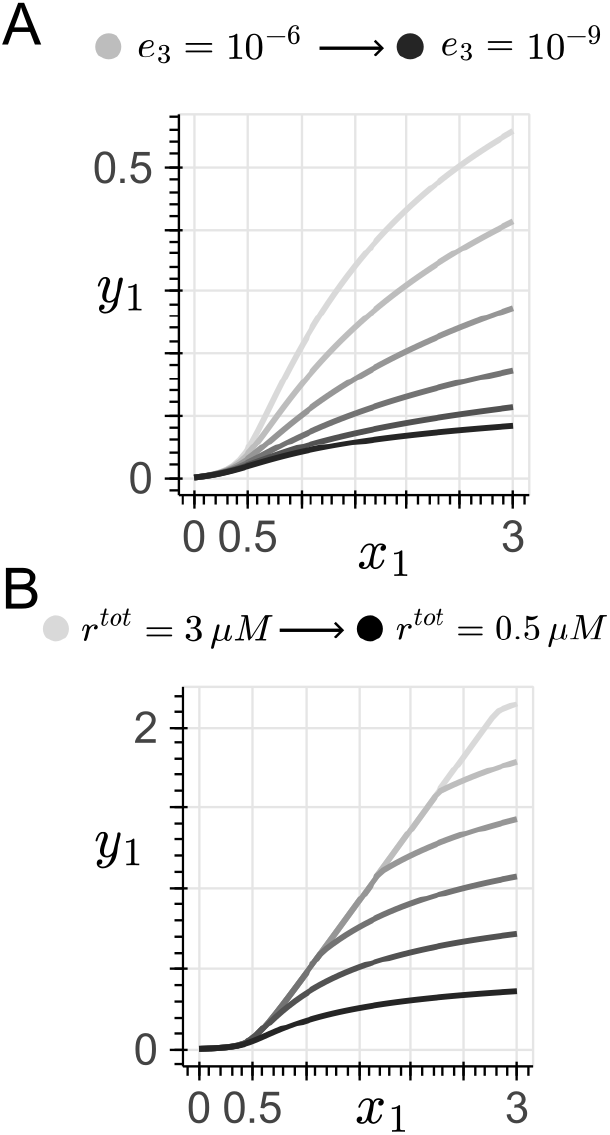
Attenuation of the output of the sequestration-based perceptron. Both panels show the steady-state activation function *ŷ*_1_ vs *x*_1_ in the homogeneous case (*e*_1_ = *e*_2_ = 10^−6^). (A) Sweep of the endogenous transcript binding affinity *e*_3_ at a fixed *x*_3_ = 1 *µM* and *r*^*tot*^ = 1 *µM* , with *e*_3_ ranging from 10^−6^ (lightest curve) to 10^−9^ (darkest curve). (B) Sweep of the total ribosome pool *r*^*tot*^ for a fixed endogenous demand (*x*_3_ = 1 *µM, e*_3_ = 100), with *r*^*tot*^ ranging from 0.5 *µM* (darkest curve) to 3 *µM* (lightest curve). In both panels, *x*_2_ = 0.5 *µM*.

## Appendix D: Closed-form expression for the ribosome allocation factor in the homogeneous regime

Denote the total circuit demand by 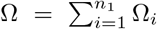. Using the relation 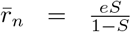, which follows from Eq. (6), and substituting into Eq. (4) yields the following quadratic equation:

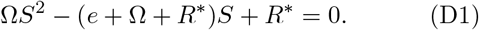

This quadratic admits a single positive root with 0 *≤ S ≤* 1. Introducing the correction factor

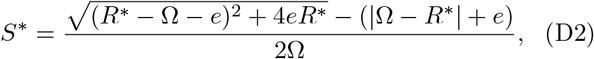

the solution can be written compactly as follows:

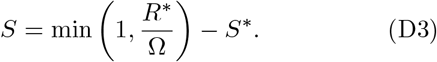

In the limit of strong binding between mRNA and ribosomes, for *e→* 0, we have that lim_*e→*0_ *S*^*\**^ = 0, and so Eq. (D3) can be written as Eq. (12).

## Appendix E: Computing the onset of resource competition in the activation function

Starting from the onset condition for resource competition, *R*^*\**^ = Ω, derived from Eq. (12), we substitute *S* = 1 and *R*^*\**^ = Ω into the ribosome conservation equation given by Eq. (4). For the homogeneous case

(*e*_1_ = *e*_2_ = *e*), this yields 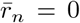. In other words, at the onset of resource competition, all free ribosomes are bound to the synthetic transcripts. Then, the endogenous allocation factor collapses to 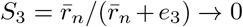 provided that *e*_3_ is finite. This regime models how the endogenous transcript dissociates from the mRNA-ribosome complexes to release ribosomes to the more strongly-binding synthetic transcripts as they approach saturation.

Substituting *S*_3_ *→* 0 into Eq. (5) results in *R*^*\**^ *→* 1, which is independent of *x*_3_ and *e*_3_. The onset of resource competition *R*^*\**^ = Ω then simplifies to Ω = 1. Expanding this term and solving for the input value *x*_1_ yields,

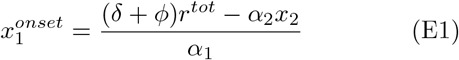

Fig. 8-A verifies numerically the approximation in Eq. (E1). For three representative values of *r*^*tot*^, with fixed *e*_1_ = *e*_2_ = 10^−6^ *≪ e*_3_ and *x*_3_ = 1 *µM* , the numerical onsets of resource competition (circles) closely match the prediction from Eq. (E1). The remaining panels of Fig. 8 illustrate the regime in which the binding affinity ordering *e*_1_ = *e*_2_ *≪ e*_3_ is relaxed. In Fig. 8-B, we sweep *e*_3_ across values that satisfy *e*_3_ *≤ e*_1,2_. In this regime, 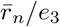 no longer collapses to zero since *e*_3_ is comparable or smaller than *e*_1,2_, so *S*_3_ exhibits a finite value, and *R*^*\**^ does not approach 1. We acknowledge that the shift of the onset approaches the limit of what has been measured experimentally for an mRNA-ribosome binding affinity, given that reproducing this regime requires the endogenous transcripts to bind even more strongly than the already strongly-binding synthetic ones, which already correspond to high-affinity RBS.

**FIG. 8.**
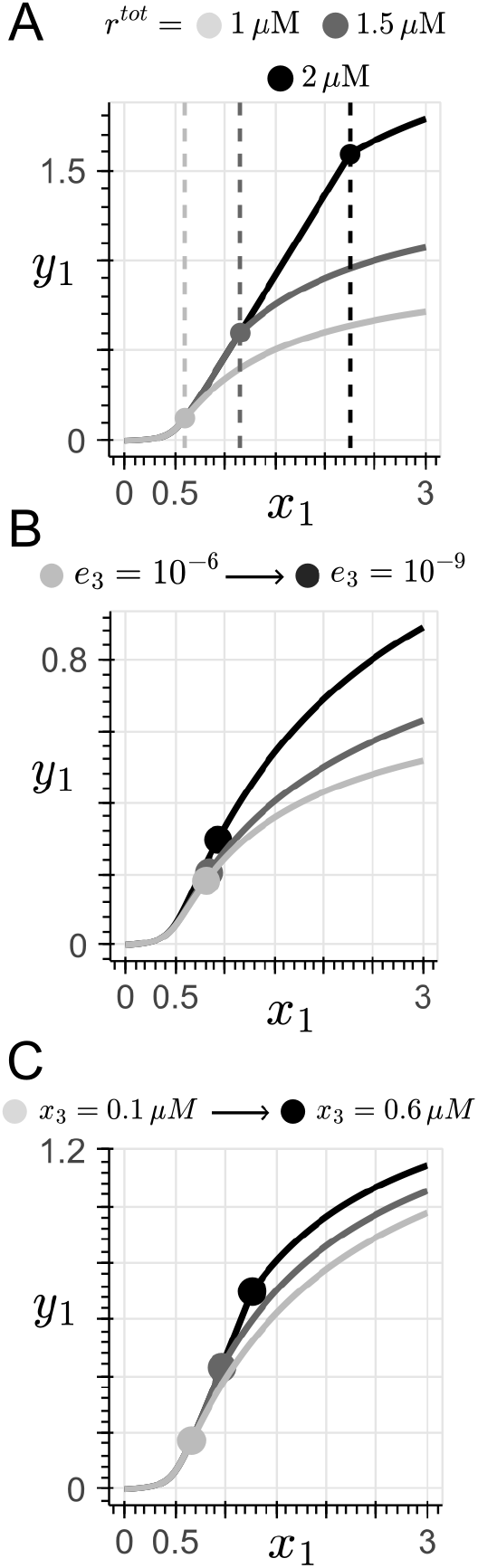
Computing the location of the onset of resource competition. All panels show the activation function 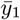 vs *x*_1_ in the homogeneous case (*e*_1_ = *e*_2_), with the numerical onset marked with a circle. (A) Verification of the analytical prediction of the onset of resource competition for the affinity ordering *e*_1,2_ *≪ e*_3_. The synthetic transcripts are held at *e*_1_ = *e*_2_ = 10^−6^, while the endogenous transcript was set at *e*_3_ = 100. The total ribosome pool is swept across *r*^*tot*^ = [1, 1.5, 2] *µM* . The dashed vertical lines indicate the closed-form prediction derived in Appendix E. (B) Relaxing the affinity ordering into a regime where *e*_3_ *≤ e*_1,2_ (i.e., the endogenous transcript binds at least as strongly as the synthetic ones). For *r*^*tot*^ = 1.6 *µM* and *e*_1_ = *e*_2_ = 10^−6^, the binding affinity *e*_3_ was swept across 10^−6^ to 10^−9^ (light to dark). (C) Onset of resource competition under matching binding affinities (*e*_1_ = *e*_2_ = *e*_3_ = 10^−6^). The input value *x*_3_ for the endogenous demand is swept across 0.1 to 0.6 *µM* , with *r*^*tot*^ = 1.6 *µM* . In all panels, *x*_2_ = 0.5 *µM*.

Consequently, a more experimentally accessible regime is one in which all transcripts share comparable binding affinities (*e*_3_ *≈ e*_1,2_), which is illustrated in Fig. 8-C. Since *S*_3_ remains finite, repeating the derivation from before (now without the *S*_3_ *→* 0 simplification), we obtain the following expression:

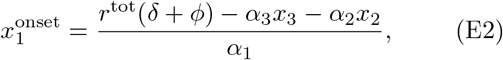

The shift to the left in the onset towards smaller input values *x*_1_ due to increases in the input level *x*_3_ of the endogenous transcript shown in Fig. 8-C is in agreement with Eq. (E2).

## Appendix F: On the location of the onset of resource competition and the bending of the decision boundary

In Section II.C.1, we showed how heterogeneity in the mRNA-ribosome binding affinity of the synthetic circuit (*e*_1_ ≠ *e*_2_) gives rise to curvature in the decision boundary. The specific set of parameters used in Fig. 4 was chosen to make the curvature easier to identify. Hence, here we analyze two complementary parameter variations to verify that the qualitative results are independent of that particular parameter regime.

Fig. 9-A and B show the free ribosome fraction 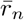 for two values of total ribosome pool. As *r*^*tot*^ decreases, the depleted (white) region expands towards the origin: a smaller ribosome pool means that free ribosomes are exhausted at smaller input values, and the onset locus of resource competition *R*^*\**^ = Ω shifts inwards accordingly. As a result, the decision boundary bends at smaller input values. Conversely, larger *r*^*tot*^ expands the abundant ribosomes region (blue), pushing the onset locus away from the origin, and delaying the bending until larger input values.

**FIG. 9.**
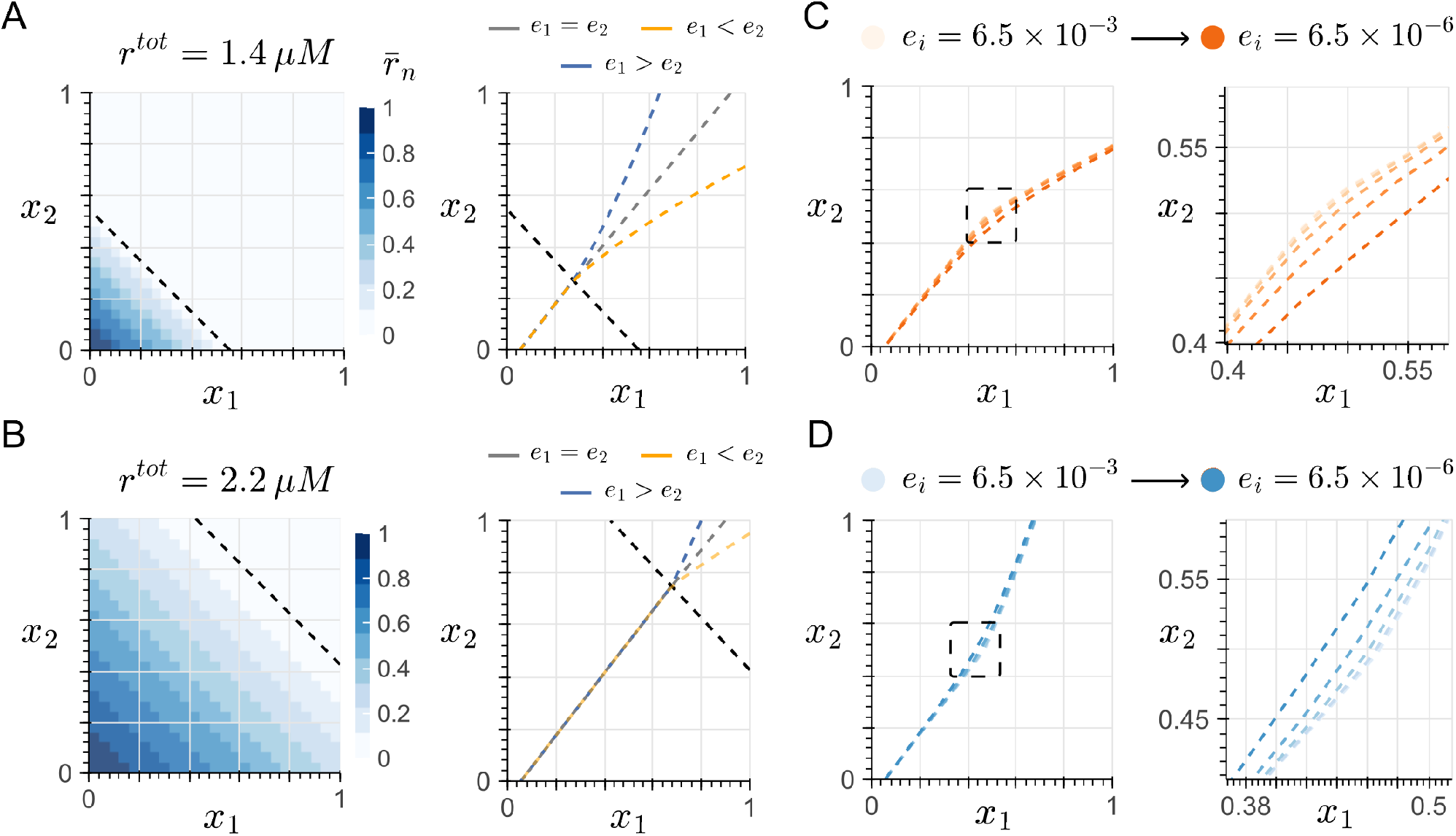
The decision boundary in the heterogeneous-affinity case across total ribosome pool (*r*^*tot*^) and bindingaffinity (*e*_*i*_) scales. (A) Left: Free ribosome fraction 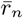 across the input space at *r*^*tot*^ = 1.4 *µM* , with the locus *R*^*\**^ = Ω overlaid as a black dashed line. Right: representative decision boundaries for the homogeneous (*e*_1_ = *e*_2_, gray) and heterogeneous (*e*_1_ < *e*_2_ in orange, and *e*_1_ > *e*_2_ in blue) affinity cases, showing the onset of bending. (B) Same as (A), but for *r*^*tot*^ = 2.2 *µM* . (C) Decision boundary for *e*_1_ < *e*_2_, sweeping from a baseline affinity *e*_*b*_ = 6.5 × 10^−3^ (lightest orange) to 6.5 × 10^−6^ (darkest orange). *e*_2_ is computed as a 3-fold increase from the baseline, while *e*_1_ = *e*_3_ = *e*_*b*_. (D) Same as (C) for *e*_1_ > *e*_2_, with the boundary bending in the opposite direction, and *e*_2_ = *e*_3_ = *e*_*b*_. In all panels, *x*_3_ = 1 *µM* , with the remaining kinetic parameters detailed in the Methods section. *r*^*tot*^ = 1.8 *µM* for Panels C and D.

Moreover, Fig. 9-C and D sweep a baseline binding affinity across three orders of magnitude. At the strongest simulated binding (*e* = 6.5 10^−6^, matching

Fig. 4), the curvature of the decision boundary is most pronounced (darker orange and blue lines in Fig. 9-C and D respectively). As the binding affinity *e* increases towards more feasible values, the curvature of the decision boundary progressively diminishes and the boundary approaches its linear shape. This effect is consistent with the framework, since stronger binding (smaller *e*_*i*_) makes the ribosome allocation factor *S*_*i*_ more sensitive to changes in 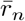, and therefore more sensitive to differences between *e*_1_ and *e*_2_. Importantly, Fig. 9-C and D validate that the direction of curvature predicted by Eq. (17) is preserved across the swept binding affinities.

## Appendix G: Mathematical models for the multi-layer perceptrons (MLPs)

Here we present the full dynamical models for the two MLP architectures shown in Fig. 5-A1 and B1. Both architectures share the same translational machinery model described by Eqs. (1)–(3) and the sequestration dynamics of Eqs. (7)–(9), applied to each node of the network. In both designs, index *i* = 3 is used to model the endogenous transcript.

An important assumption in both architectures is that the output proteins of the hidden-layer nodes (Node 1 and Node 2) drive transcription in the output node (Node 3) through a linear activation regime. Specifically, the steady-state protein concentration from a hiddenlayer node enters as the input signal in the transcription term of the downstream node (i.e., *α*_*i*_*y*_*j*_ replaces *α*_*i*_*x*_1_ in Eq. (1)). This assumes that the connections between nodes operate in a regime where protein concentration is proportional to transcriptional output, avoiding additional nonlinearities that would arise from saturating transcription factor-promoter interactions or cooperative binding. Incorporating such nonlinear activation between layers is left for future work.

### 1. Architecture A: closed decision boundary (Fig. 5-A1)

This architecture consists of two hidden-layer nodes and one output node, with a total of eight ribosome– mRNA complexes, including the endogenous complex, competing for the shared pool. The free ribosome concentration is given by

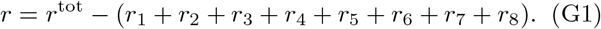

#### a. Endogenous demand (i = 3)

The endogenous transcript follows Eqs. (1)–(3) with input *x*_3_.

#### b. Node 1 (i = {1, 2})

Node 1 receives both external inputs *x*_1_ and *x*_2_. Transcripts *M*_1_ and *M*_2_ follow Eqs. (1)–(2) with inputs *x*_1_ and *x*_2_, respectively. Their translated proteins *y*_1_ and *y*_2_ undergo molecular sequestration:

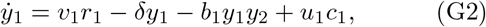

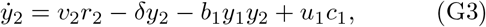

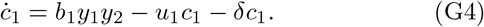

The output of Node 1 is the free protein 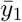.

#### c. Node 2 (i = {4, 5, 6})

Node 2 receives inputs *x*_1_ and *x*_2_ through transcripts *M*_4_ and *M*_5_, which follow Eqs. (1)–(2). Both produce the same protein *y*_4_, so their translational contributions are summed. A third transcript *M*_6_, driven by the bias input *x*_0_, produces the sequestering protein *y*_5_:

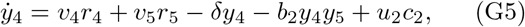

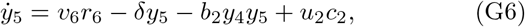

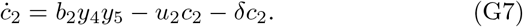

The output of Node 2 is the free protein 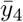.

#### d. Output node (i = {7, 8})

The output node receives the steady-state outputs of Node 1 and Node 2 as transcriptional inputs. Transcript *M*_7_ is driven by 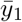 and transcript *M*_8_ by 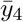, both following Eqs. (1)–(2). Their translated proteins undergo sequestration:

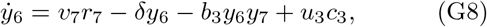

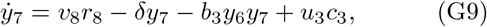

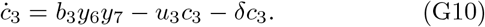

The output of this architecture is the complex 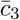.

#### 2. Architecture B: non-convex decision boundary (Fig. 5-B1)

This architecture consists of two hidden-layer nodes and one output node, with a total of ten ribosome– mRNA complexes, including the endogenous complex, competing for the shared pool. The free ribosome concentration is given by

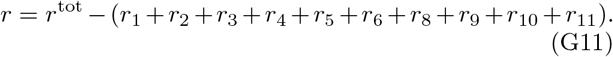

#### a. Endogenous demand (i = 3)

As in Architecture A, the endogenous transcript follows Eqs. (1)–(3) with input *x*_3_.

#### b. Node 1 (i = 1, 2, 4)

Node 1 receives both external inputs. Transcript *M*_1_ is driven by *x*_1_ and *M*_2_ by *x*_2_, producing proteins *y*_1_ and *y*_2_, respectively. Transcript *M*_4_, driven by *x*_0_, produces the sequestering protein *y*_2_. Transcripts *M*_2_ and *M*_4_ both contribute to *y*_2_:

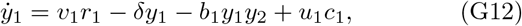

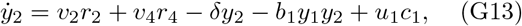

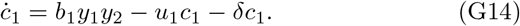

The output of Node 1 is the free protein 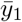.

#### c. Node 2 (i = {5, 6, 8})

Node 2 has the opposite sign assignment. Transcript *M*_5_ is driven by *x*_2_ and *M*_6_ by *x*_1_, producing proteins *y*_4_ and *y*_5_, respectively. Transcript *M*_8_, driven by *x*_0_, produces the sequestering protein *y*_5_. Transcripts *M*_6_ and *M*_8_ both contribute to *y*_5_:

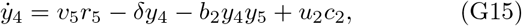

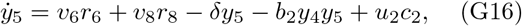

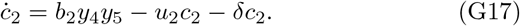

The output of Node 2 is the free protein 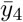.

#### d. Output node (i = {9, 10, 11})

The output node receives the steady-state outputs of Node 1 and Node 2, along with a bias. Transcripts *M*_9_ and *M*_11_ are driven by 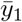 and 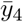, respectively, and both produce protein *y*_6_. Transcript *M*_10_, driven by *x*_0_, produces the sequestering protein *y*_7_:

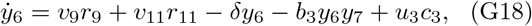

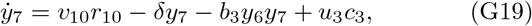

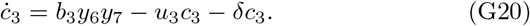

The output of this architecture is the free protein 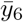.

## Appendix H: Analysis of the decision boundary of a single perceptron with an inhibitory bias

Consider Eqs. (1)–(3) for *i* = 1, 2, 3, where indices 1 and 2 correspond to two mRNA molecules *M*_1_ and *M*_2_ whose transcription is driven by the inputs *x*_1_ and *x*_2_, respectively, and index 3 corresponds to a third mRNA *M*_3_ whose transcription is driven by the bias input *x*_0_. Both *M*_1_ and *M*_2_ are translated into the same protein *Y*_1_, while *M*_3_ produces the protein *Y*_2_. We model the dynamics of this sequestration reaction using the following ODEs:

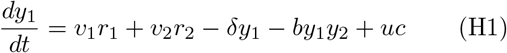

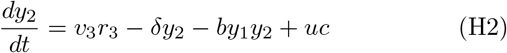

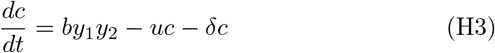

Setting Eq. (H3) to zero and solving the steady state of Eqs. (H1) and (H2) in terms of 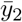, we obtain

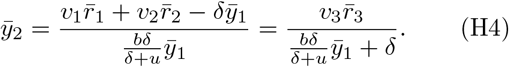

Here 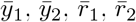 and 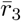 denote the steady-state values of *y*_1_(*t*), *y*_2_(*t*), *r*_1_(*t*), *r*_2_(*t*), and *r*_3_(*t*), respectively. Substituting 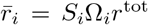 from Eq. (A3) and defining *ξ* = (*δ* + *u*)*/b*, the decision boundary can be written compactly as

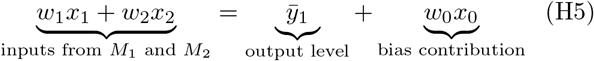

where, considering the ribosome allocation factor from Eq. (6), the effective weights are

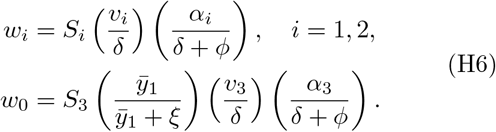

Note that Eq. (H5) highlights two effects of ribosome competition. First, the bias term *w*_0_*x*_0_ acts as a fixed threshold, so any change in *w*_0_, whether from varying *M*_3_ or a global shift in 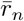, translates the boundary without changing its slope *Q*. Second, differences in ribosomebinding affinities between *M*_1_ and *M*_2_ have a minimal effect on the curvature of the decision boundary. Since *x*_1_ and *x*_2_ appear on the same side of the boundary, increases in one input along the boundary must be compensated by decreases in the other, keeping total ribosome demand approximately constant. Consequently, 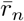 is nearly uniform along the boundary, *Q*’s variation is negligible, and the boundary slants rather than bends. In contrast, in the previously analyzed perceptron, where inputs are on opposite sides of the boundary, 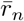 varies substantially, producing visible curvature.

## Appendix I: The decision boundary under growth-mediated feedback

### 1. Steady-state derivations under *δ → λ* **and a variable** *r*^*tot*^

Following the steps of Appendix A with these substitutions, the steady-state ribosome-mRNA complex can be written as:

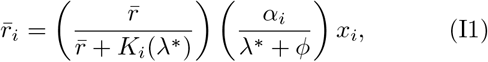

where

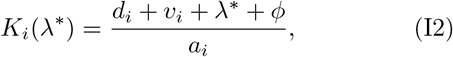

Normalizing by *r*^*tot,\**^ we define the growth-dependent dimensionless parameters

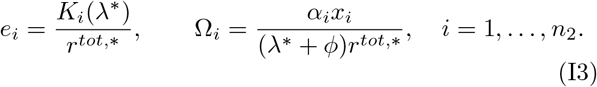

Note that the ribosome conservation relation described by Eq. (4) and the definition of the ribosome allocation factor *S*_*i*_ in Eq. (6) conserve their form under these substitutions, establishing the growth-feedback invariance of this framework.

#### 2. The slope of the decision boundary

The form of the slope of the decision boundary described by Eq. (14) is preserved under growth-mediated feedback, with *e*_*i*_ and 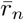 now evaluated at (*λ*^*\**^, *r*^*tot,\**^). In the homogeneous-affinity case (*e*_1_ = *e*_2_), the factor 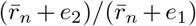 reduces to unity for any *λ*^*\**^ and *r*^*tot,\**^, so the slope is fixed only by the transcription and translation rates.

Moreover, to analyze the heterogeneous case (*e*_1_ ≠ *e*_2_), from Eq. (15), we take the derivative of the slope *q* = *S*_1_*/S*_2_ with respect to an input *x*_*i*_. With 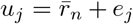 so that *q* = *u*_2_*/u*_1_, and applying the chain rule, we obtain the following equation:

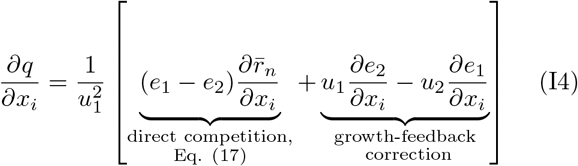

Since the factor 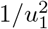 is strictly positive, the sign of the expression is determined by the sign within the brackets. Analyzing the first term, given that increasing the expression of any synthetic circuit consumes ribosomes, and therefore reduces the free, available pool, we get 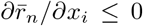, which can be obtained otherwise by differentiating the ribosome mass conservation relation in Eq. (4) with respect to *x*_*i*_. Therefore, we obtained the same dependency on the ordering between *e*_1_ and *e*_2_ from Eq. (17).

On the other hand, analyzing the second term is less straightforward, since *e*_*j*_ = *K*_*j*_(*λ*^*\**^)*/r*^*tot,\**^. In other words, *e*_*j*_ depends on both *λ*^*\**^ and *r*^*tot,\**^, resulting in

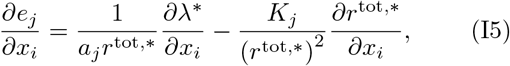

where *a*_*j*_ is the ribosome-mRNA association constant for the *j*-th transcript, and *K*_*j*_(*λ*^*\**^) = (*d*_*j*_ + *v*_*j*_ + *λ*^*\**^ + *ϕ*)*/a*_*j*_, both introduced in Appendix A. Since all rate constants and the total pool of ribosomes are nonnegative, the sign of Eq. (I5) is determined by the derivatives it contains. First, since increasing gene expression increases the translational burden and therefore reduces growth [14], *∂λ*^*\**^*/∂x*_*i*_ *≤* 0. Second, the increase in gene expression reduces the ribosomal fraction of the total proteome [14, 19, 26], so *∂r*^tot,*^*/∂x*_*i*_ *≤* 0. Overall, the two terms in Eq. (I5) have opposite signs, and the sign and magnitude of *∂e*_*j*_*/∂x*_*i*_ therefore depend on the relative strength of each effect, which in turn depends on the specific form of Ψ.

In summary, under growth-mediated feedback, the direction of bending of the decision boundary depends not only on the ordering of *e*_*j*_, but also on host physiology, represented by Ψ. Nevertheless, since the values of *λ*^*\**^ and *r*^*tot,\**^ are within a bounded range for any specific host [29], the resulting decision boundary is constrained to a region in the input space. The design of a classifier around this uncertainty remains for future work.

#### 3. Bending of the onset locus of resource competition

Whether the onset locus *R*^*\**^ = Ω bends in the input space depends on the structure of Ψ. In particular, for phenomenological growth laws in which growth is dependent only on translational burden, such as those proposed by [14], the onset locus remains linear. That is, for growth laws of the form

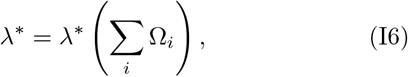

the onset condition fixes Σ_*i*_ Ω_*i*_ to a specific value, and therefore *λ*^*\**^ and *r*^*tot,\**^ are constant along the locus, which reduces to

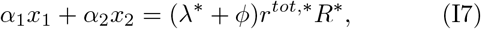

evaluated at the onset. Bending of this locus requires the growth law to resolve burden between individual transcripts, for instance, by detailing differences in protein length, codon usage, or expression demand [19, 28].

## References

[1] A. Adelaja, B. Taylor, K. M. Sheu, Y. Liu, S. Luecke, and A. Hoffmann, Six distinct nfb signaling codons convey discrete information to distinguish stimuli and enable appropriate macrophage responses, Immunity 54, 916 (2021).

[2] P. Li and M. B. Elowitz, Communication codes in developmental signaling pathways, Development 146, 10.1242/dev.170977 (2019).

[3] A. A. Granados, N. Kanrar, and M. B. Elowitz, Combinatorial expression motifs in signaling pathways, Cell Genomics 4, 100463 (2024).

[4] F. Britto Bisso, R. Aguilar, D. Shree, Y. Zhu, M. Espinoza, B. Diaz, and C. Cuba Samaniego, Pattern recognition in living cells through the lens of machine learning, Open Biology 15, 10.1098/rsob.240377 (2025).

[5] L. Qian, E. Winfree, and J. Bruck, Neural network computation with dna strand displacement cascades, Nature 475, 368 (2011).

[6] K. M. Cherry and L. Qian, Scaling up molecular pattern recognition with dna-based winner-take-all neural networks, Nature 559, 370–376 (2018).

[7] K. R. Rodriguez, N. Sarraf, and L. Qian, A loser-take-all dna circuit, ACS Synthetic Biology 10, 2878–2885 (2021).

[8] R. Lopez, R. Wang, and G. Seelig, A molecular multi-gene classifier for disease diagnostics, Nature Chemistry 10, 746–754 (2018).

[9] E. Nakamura, F. B. Bisso, I. Gispert, H. Moghimianavval, S. Okuda, A. Przybyszewska-Podstawka, S. Chisholm, V. S. Hemanth Kumar, V. P. Medina, M.-R. Wu, and C. C. Samaniego, Sequestration-based neural networks that operate out of equilibrium 10.1101/2025.05.16.654484 (2025).

[10] A. Moorman, Machine learning inspired Synthetic Biology: Neuromorphic computing in mammalian cells, Ph.D. thesis (2020).

[11] Z. Chen, J. M. Linton, S. Xia, X. Fan, D. Yu, J. Wang, R. Zhu, and M. B. Elowitz, A synthetic protein-level neural network in mammalian cells, Science 386, 1243–1250 (2024).

[12] M. Scott and T. Hwa, Shaping bacterial gene expression by physiological and proteome allocation constraints, Nature Reviews Microbiology 21, 327–342 (2022).

[13] Y. Qian, H.-H. Huang, J. I. Jiménez, and D. Del Vecchio, Resource competition shapes the response of genetic circuits, ACS Synthetic Biology 6, 1263–1272 (2017).

[14] M. Scott, C. W. Gunderson, E. M. Mateescu, Z. Zhang, and T. Hwa, Interdependence of cell growth and gene expression: Origins and consequences, Science 330, 1099–1102 (2010).

[15] E.-M. Nikolados, A. Y. Weiße, F. Ceroni, and D. A. Oyarzún, Growth defects and loss-of-function in synthetic gene circuits, ACS Synthetic Biology 8, 1231–1240 (2019).

[16] A. Gyorgy, J. I. Jiménez, J. Yazbek, H.-H. Huang, H. Chung, R. Weiss, and D. Del Vecchio, Isocost lines describe the cellular economy of genetic circuits, Biophysical Journal 109, 639–646 (2015).

[17] T. E. Gorochowski, I. Avcilar-Kucukgoze, R. A. L. Bovenberg, J. A. Roubos, and Z. Ignatova, A minimal model of ribosome allocation dynamics captures trade-offs in expression between endogenous and synthetic genes, ACS Synthetic Biology 5, 710–720 (2016).

[18] F. Ceroni, R. Algar, G.-B. Stan, and T. Ellis, Quantifying cellular capacity identifies gene expression designs with reduced burden, Nature Methods 12, 415–418 (2015).

[19] K. Sechkar, H. Steel, G. Perrino, and G.-B. Stan, A coarse-grained bacterial cell model for resource-aware analysis and design of synthetic gene circuits, Nature Communications 15, 10.1038/s41467-024-46410-9 (2024).

[20] C. D. McBride and D. Del Vecchio, Predicting composition of genetic circuits with resource competition: demand and sensitivity, ACS Synthetic Biology 10, 3330 (2021).

[21] A. Moorman, C. C. Samaniego, C. Maley, and R. Weiss, A dynamical biomolecular neural network, in 2019 IEEE 58th Conference on Decision and Control (CDC) (IEEE, 2019).

[22] C. C. Samaniego, Y. Qian, K. Carleton, and E. Franco, Building subtraction operators and controllers via molecular sequestration, IEEE Control Systems Letters 7, 3361–3366 (2023).

[23] F. Britto Bisso, D. Shree, Y. Zhu, and C. Cuba Samaniego, Design principles of neuromorphic computing using genetic circuits 10.64898/2025.12.01.691482 (2025).

[24] H. M. Salis, E. A. Mirsky, and C. A. Voigt, Automated design of synthetic ribosome binding sites to control protein expression, Nature Biotechnology 27, 946–950 (2009).

[25] C. C. Samaniego, E. Wallace, F. Blanchini, E. Franco, and G. Giordano, Neural networks built from enzymatic reactions can operate as linear and nonlinear classifiers, in 2024 IEEE 63rd Conference on Decision and Control (CDC) (IEEE, 2024) p. 6292–6297.

[26] N. A. Vaccari, D. Zevallos-Aliaga, T. Peeters, and D. G. Guerra, Biosensor characterization: formal methods from the perspective of proteome fractions, Synthetic Biology 10, 10.1093/synbio/ysaf002 (2025).

[27] J. C. Kratz, H. Wang, F. Si, and S. Banerjee, Multitimescale adaptation and emergent learning in single bacterial cells, PRX Life 4, 10.1103/5zbg-8vll (2026).

[28] G. Chure and J. Cremer, An optimal regulation of fluxes dictates microbial growth in and out of steady state, eLife 12, 10.7554/elife.84878 (2023).

[29] M. Mori, Z. Zhang, A. Banaei-Esfahani, J. Lalanne, H. Okano, B. C. Collins, A. Schmidt, O. T. Schubert, D. Lee, G. Li, R. Aebersold, T. Hwa, and C. Ludwig, From coarse to fine: the absolute escherichia coli proteome under diverse growth conditions, Molecular Systems Biology 17, 10.15252/msb.20209536 (2021).

[30] C. C. Samaniego, A. Moorman, G. Giordano, and E. Franco, Signaling-based neural networks for cellular computation, in 2021 American Control Conference (ACC) (IEEE, 2021) pp. 1883–1890.

[31] A. Schmidt, K. Kochanowski, S. Vedelaar, E. Ahrńe, B. Volkmer, L. Callipo, K. Knoops, M. Bauer, R. Aebersold, and M. Heinemann, The quantitative and condition-dependent escherichia coli proteome, Nature Biotechnology 34, 104–110 (2016).

[32] R. D. Jones, Y. Qian, V. Siciliano, B. DiAndreth, J. Huh, R. Weiss, and D. Del Vecchio, An endoribonuclease-based feedforward controller for decoupling resource-limited genetic modules in mammalian cells, Nature Communications 11, 10.1038/s41467-020-19126-9 (2020).

[33] R. Zhang, R. Zhang, and X.-J. Tian, Integrating recombinase-based feedback and feedforward control for optimal resource decoupling, Nucleic Acids Research 53, 10.1093/nar/gkaf913 (2025).

[34] T. Frei, C.-H. Chang, M. Filo, A. Arampatzis, and M. Khammash, A genetic mammalian proportional–integral feedback control circuit for robust and precise gene regulation, Proceedings of the National Academy of Sciences 119, 10.1073/pnas.2122132119 (2022).

[35] F. Ceroni, A. Boo, S. Furini, T. E. Gorochowski, O. Borkowski, Y. N. Ladak, G. A. Nicholson, D. P. Storch, R. T. S. King, A. M. H. K. Reddington, T. Wei, L. S. Vind, J. Casanovas, T. Dang, G. Giordano, Y. Liang, and T. Ellis, Burden-driven feedback control of gene expression, Nature Methods 15, 387 (2018).

[36] A. Stone, J. Ryan, X. Tang, and X.-J. Tian, Negatively competitive incoherent feedforward loops mitigate winner-take-all resource competition, ACS Synthetic Biology 11, 3986–3995 (2022).

[37] R. D. Jones, Y. Qian, K. Ilia, B. Wang, M. T. Laub, D. Del Vecchio, and R. Weiss, Robust and tunable signal processing in mammalian cells via engineered covalent modification cycles, Nature Communications 13, 10.1038/s41467-022-29338-w (2022).

[38] H.-H. Huang, Y. Qian, and D. Del Vecchio, A quasiintegral controller for adaptation of genetic modules to variable ribosome demand, Nature Communications 9, 10.1038/s41467-018-07899-z (2018).

[39] S. Chakravarty, R. Guttal, R. Zhang, and X.-J. Tian, Mitigating winner-take-all resource competition through antithetic control mechanism, ACS Synthetic Biology 13, 4050–4060 (2024).

[40] Y. Qian and D. Del Vecchio, Mitigation of ribosome competition through distributed srna feedback, in 2016 IEEE 55th Conference on Decision and Control (CDC) (IEEE, 2016) p. 758–763.

[41] F. Nóbel and J. Picó, Resources allocation explains the differential roles of rbs and promoter strengths in cell mass distribution and optimal protein expression productivity 10.1101/2020.11.19.390583 (2020).

[42] O. Rackham and J. W. Chin, A network of orthogonal ribosome·mrna pairs, Nature Chemical Biology 1, 159–166 (2005).

[43] N. A. Aleksashin, T. Szal, A. E. d’Aquino, M. C. Jewett, N. Vázquez-Laslop, and A. S. Mankin, A fully orthogonal system for protein synthesis in bacterial cells, Nature Communications 11, 10.1038/s41467-020-15756-1 (2020).

[44] Y. Duan, X. Zhang, W. Zhai, J. Zhang, X. Zhang, G. Xu, H. Li, Z. Deng, J. Shi, and Z. Xu, Deciphering the rules of ribosome binding site differentiation in context dependence, ACS Synthetic Biology 11, 2726–2740 (2022).

[45] D. Omotajo, T. Tate, H. Cho, and M. Choudhary, Distribution and diversity of ribosome binding sites in prokaryotic genomes, BMC Genomics 16, 10.1186/s12864-015-1808-6 (2015).

[46] N. Senda, T. Enomoto, K. Kihara, N. Yamashiro, N. Takagi, D. Kiga, and H. Nishida, Development of an expression-tunable multiple protein synthesis system in cell-free reactions using t7-promoter-variant series, Synthetic Biology 7, 10.1093/synbio/ysac029 (2022).

[47] R. Milo, P. Jorgensen, U. Moran, G. Weber, and M. Springer, Bionumbers—the database of key numbers in molecular and cell biology, Nucleic Acids Research 38, D750 (2010).

[48] S. Kosuri, D. B. Goodman, G. Cambray, V. K. Mutalik, Y. Gao, A. P. Arkin, D. Endy, and G. M. Church, Composability of regulatory sequences controlling transcription and translation in escherichia coli, Proceedings of the National Academy of Sciences 110, 14024–14029 (2013).

[49] H. Moghimianavval, I. Gispert, S. R. Castillo, O. B. Corning, A. P. Liu, and C. Cuba Samaniego, Engineering sequestration-based biomolecular classifiers with shared resources, ACS Synthetic Biology 13, 3231 (2024).

[50] G.-W. Li, D. Burkhardt, C. Gross, and J. S. Weissman, Quantifying absolute protein synthesis rates reveals principles underlying allocation of cellular resources, Cell 157, 624–635 (2014).

[51] D. Del Vecchio, Modularity, context-dependence, and insulation in engineered biological circuits, Trends in Biotechnology 33, 111–119 (2015).

[52] G. Cambray, J. C. Guimaraes, and A. P. Arkin, Evaluation of 244, 000 synthetic sequences reveals design principles to optimize translation in escherichia coli, Nature Biotechnology 36, 1005–1015 (2018).

[53] V. K. Mutalik, J. C. Guimaraes, G. Cambray, C. Lam, M. J. Christoffersen, Q.-A. Mai, A. B. Tran, M. Paull, J. D. Keasling, A. P. Arkin, and D. Endy, Precise and reliable gene expression via standard transcription and translation initiation elements, Nature Methods 10, 354–360 (2013).

[54] Z. Zhao, X. Liu, W. Zhang, Y. Yang, X. Dai, and Z. Bai, Construction of genetic parts from the corynebacterium glutamicum genome with high expression activities, Biotechnology Letters 38, 2119–2126 (2016).

[55] S. H. Jang, J. W. Cha, N. S. Han, and K. J. Jeong, Development of bicistronic expression system for the enhanced and reliable production of recombinant proteins in leuconostoc citreum, Scientific Reports 8, 10.1038/s41598-018-27091-z (2018).

